# Profound geroprotection from brief rapamycin treatment in early adulthood by persistently increased intestinal autophagy

**DOI:** 10.1101/2022.04.20.488884

**Authors:** Paula Juricic, Yu-Xuan Lu, Thomas Leech, Lisa F. Drews, Jonathan Paulitz, Jiongming Lu, Tobias Nespital, Sina Azami, Jennifer C. Regan, Emilie Funk, Jenny Fröhlich, Sebastian Grönke, Linda Partridge

## Abstract

The licensed drug rapamycin has potential to be repurposed for geroprotection. A key challenge is to avoid adverse side-effects from continuous dosing regimes. Here we show a profound memory effect of brief, early rapamycin treatment of adults, which extended lifespan in *Drosophila* to the same degree as lifelong dosing. Lasting memory of earlier rapamycin treatment was mediated by elevated autophagy in enterocytes of the gut, accompanied by increased levels of intestinal lysosomal alpha-mannosidase V (LManV), proteins involved in branched-chain amino acid metabolism and lysozyme levels, and by maintained structure and function of the ageing intestine. Brief elevation of autophagy in early adulthood itself induced a long-term increase in autophagy. In mice, a short-term, 3-month treatment in early adulthood also induced a memory effect, with maintenance similar to that seen with chronic treatment, of lysozyme distribution, Man2B1 level in intestinal crypts, Paneth cell architecture and gut barrier function, even 6 months after rapamycin was withdrawn. The geroprotective effects of chronic rapamycin treatment can thus be obtained with a brief pulse of the drug in early adulthood.

## Main

The macrolide drug rapamycin inhibits TORC1 activity and can extend lifespan in model organisms, including mice^1-3^. In mice rapamycin can delay several age-related diseases, such as cognitive decline^4^, spontaneous tumours^5^, and cardiovascular^6,7^ and immune dysfunction^8^. However, chronic rapamycin administration can cause adverse effects, even with low doses^9,10^. Shortening treatment could potentially reduce negative effects. Short-term treatment in late life can extend lifespan in mice^3,11,12^ and enhance immune response in elderly humans^13,14^. However, it is unknown whether the effects of late-life treatment are comparable to those of lifelong drug exposure, or whether brief treatment at younger ages is sufficient to gain the benefits of the chronic treatment.

To assess the efficacy of late-onset rapamycin treatment, we treated *Drosophila* at different ages and for varying durations. Treatments starting later in life, on day 30 or day 45, extended lifespan, consistent with previous findings in mice^3,11,12^, but less than did lifelong treatment (Fig. 1a-b, and Extended Data Table 1 and 2). Very late-onset rapamycin treatment from day 60, when survival is already decreased to ∼80%, did not increase lifespan (Fig. 1c and Extended Data Table 1). Thus, later onset rapamycin treatment produced progressively smaller extensions of lifespan.

**Fig. 1.**
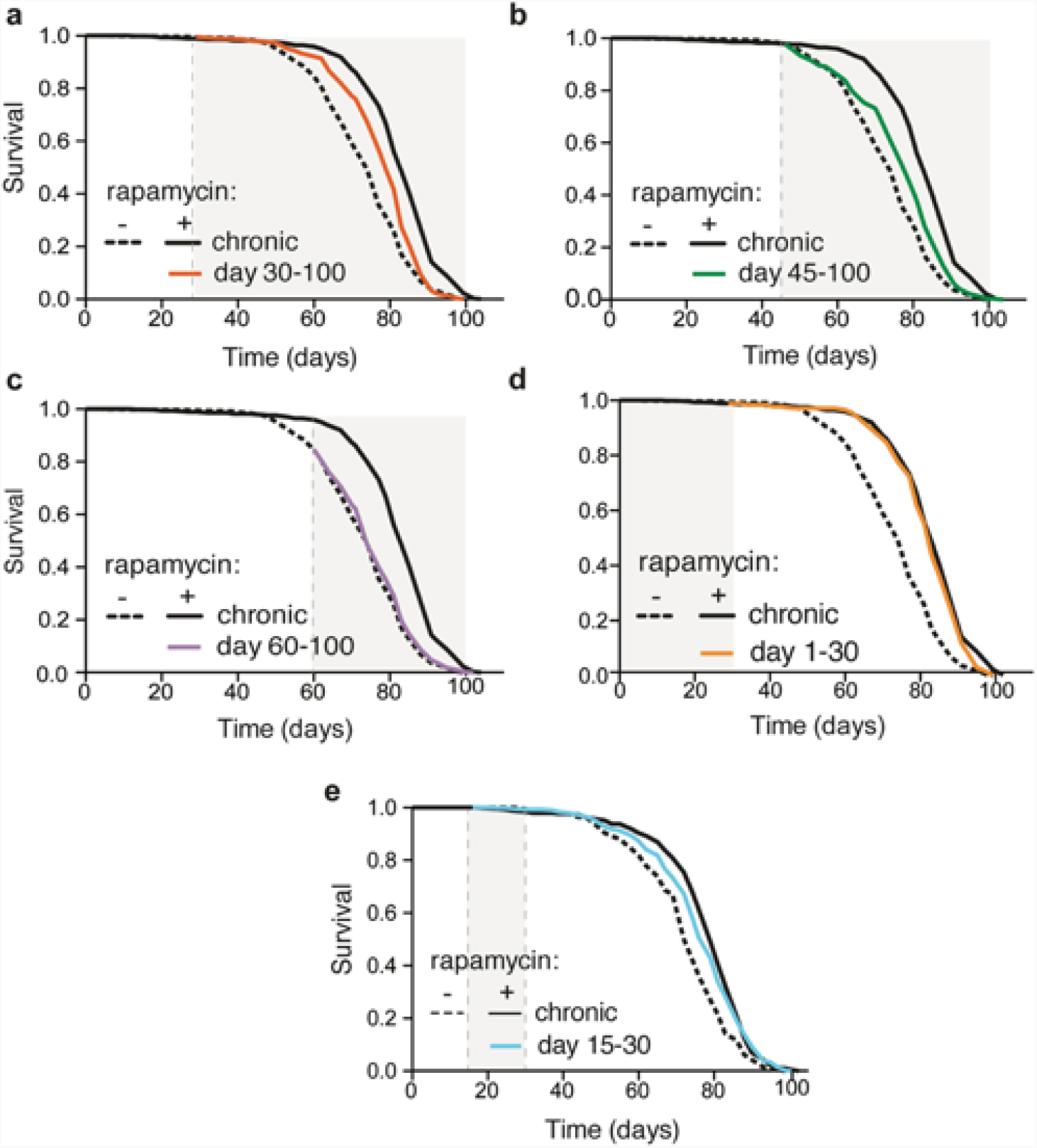
Lifespan response to rapamycin treatment declines with the age of onset of the treatment. **a**, Rapamycin treatment started on day 30 extended lifespan (p=2.13 × 10^−06^) to a lesser degree than did lifelong treatment (p=1.04 × 10^−13^, see also Extended Data Table 2). **b-c**, Rapamycin treatment started on day 45 modestly extended lifespan (**b**, p=0.0003, see also Extended Data Table 1) whereas treatment started on day 60 (**c**) had no lifespan-extending effect (p=0.256, see also Extended Data Table 1). d, Rapamycin treatment from day 1-30 extended lifespan (p=2.13 × 10^−06^) as much as did chronic treatment (p=0.09, see also Extended Data Table 2). **e**, Treatment from days 15-30 extended lifespan slightly less than did chronic treatment (d15-30 vs. control: p=7.58 × 10^−07^; d15-30 vs. chronic rapamycin: p=0.19, see also Extended Data Table 3). Note that the experiments in Extended Data Fig. 1a-d were run in parallel, hence the lifespan data of the control flies is the same. Experiments in Fig. 1e and 2a were run in parallel, therefore lifespan data of the control flies is the same. N=400 flies per condition.

In sharp contrast, rapamycin treatment instigated early in adulthood on day 3 following eclosion and 2 days of mating (termed “day 1”), for just 30 days, extended lifespan as much as did lifelong dosing (Fig. 1d and Extended Data Table 2). Treatment from day 15-30 increased lifespan, but less than did chronic treatment (Fig. 1e and Extended Data Table 3). Remarkably, rapamycin in only the first 15 days of adult life recapitulated the full lifespan extension achieved by chronic treatment (Fig. 2a and Extended Data Table 3), a phenomenon we termed ‘rapamycin memory’.

**Fig. 2:**
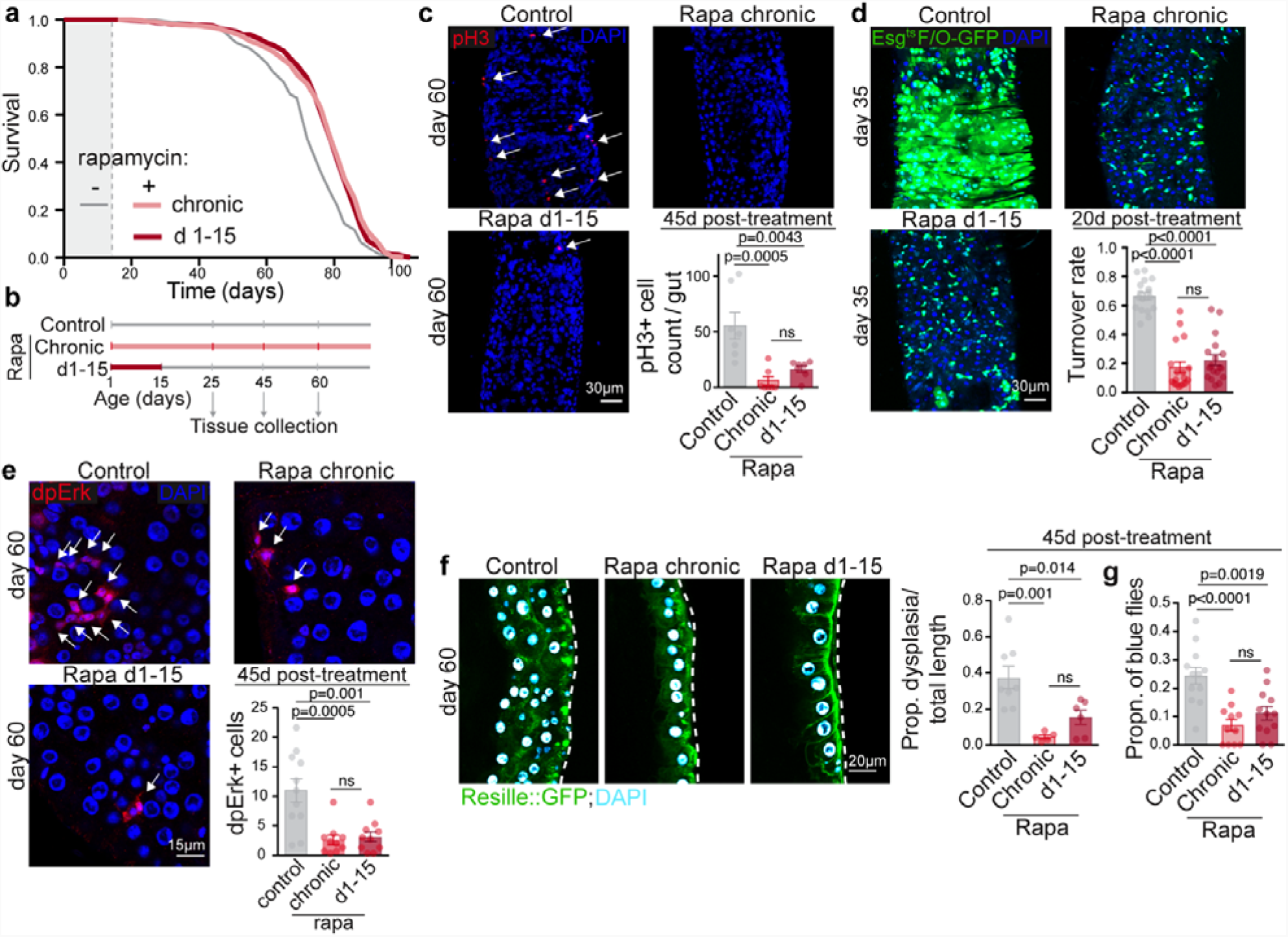
Brief rapamycin treatment early in adulthood extends lifespan and preserves intestinal function as much as does chronic treatment. **a**, Lifespan of flies chronically or in days 1-15 treated with rapamycin (n=400 per condition, see also Extended Data Table 3). **b**, Experimental design. **c**, The number of pH3+ cells (arrows) in the gut 45 days after the short-term rapamycin treatment was withdrawn (n=7-8). d, Midgut turnover rate, as assessed with the esg^ts^F/O system 20 days post-treatment (n=15-18). **e**, The number of dpErk+ cells 45-days post-rapamycin treatment (n=10-11). **f**, Intestinal dysplasia in gut R2 region of flies carrying epithelial marker Resille-GFP 45-days after short-term rapamycin treatment was terminated (n=6-8). **g**, Intestinal barrier function in flies treated with rapamycin chronically or in days 1-15. Data are mean ± s.e.m. One-way ANOVA, Bonferroni’s post-test.

Rapamycin increases lifespan mainly in female *Drosophila* ^2^. The number of dividing intestinal stem cells (ISCs) increases with age in female flies, to restore damaged parts of the intestinal epithelium, driving intestinal dysplasia later in life^15^. Thus, we hypothesized that short-term rapamycin might permanently alter ISC activity. As previously reported^16^, chronic rapamycin treatment reduced pH3+ cell number (Fig. 2c), a marker for dividing cells^17^. Strikingly, the number of pH3+ cells of flies treated with rapamycin only during days 1-15 remained as low as in flies treated chronically, even 10, 30 and 45 days post-treatment (Fig. 2b-c; Extended Data Fig. 1a-b). Mass spectrometry confirmed that rapamycin concentration was reduced to the level of control flies 10-days after rapamycin treatment on days 1-15 was ended (Extended Data Fig. 1c). The ISCs thus remained fully quiescent long after rapamycin had been cleared.

We next assessed the turnover rate of the intestinal epithelium using the *esg*^*ts*^ *F/O system (esg-Gal4; tubGal80*^*ts*^ *Act>CD2>Gal4 UAS-Flp UAS-GFP)*^18^, where activation by a temperature shift to 29°C marks ISCs and their progenitor cells with GFP. Under standard conditions, the epithelial turnover rate in *Drosophila* is 14 days. Temperature increase shortens lifespan, so we measured turnover rate 10 and 20 days post-treatment. Most of the control midgut epithelium was replaced by GFP positive cells after 10 (Extended Data Fig. 1d) and 20 days (Fig. 2d) of system activation. Chronic and day 1-15 rapamycin treatment reduced the number of GFP positive cells 10 and 20 days after the switch to the same extent (Fig. 2d; Extended Data Fig. 1d). Brief, early rapamycin exposure thus reduced turnover of the intestinal epithelium as much as chronic treatment, and the cells previously treated with rapamycin remained in the gut until advanced age.

Staining with diphosphorylated Erk (dpErk), a specific readout for signal that damaged and apoptotic enterocytes send to the ISCs for replacement^19^, revealed that short-term rapamycin treatment reduced the number of apoptotic, dpErk positive cells as much as did chronic treatment (Fig. 2e), suggesting increased enterocyte health. We therefore assessed if intestinal pathologies were reduced. Histology using the epithelial marker *Resille-GFP* revealed that dysplastic regions were widespread throughout the gut of ageing control flies (Fig. 2f). Flies treated chronically with rapamycin had significantly fewer dysplastic lesions at day 60. Interestingly, proportion of dysplastic regions remained reduced 45 days after short-term rapamycin treatment was withdrawn, to the same degree as seen with chronic treatment (Fig. 2f). Since lifespan is directly linked to gut barrier function, and loss of septate-junction proteins disrupts gut integrity^20^, we measured the effect of brief rapamycin treatment on gut barrier function. Intestinal integrity, as measured by a blue dye leakage assay, was preserved by rapamycin treatment, and remained fully protected even 45 days after rapamycin was withdrawn (Fig. 2g). Taken together, these results indicate that brief, early-life rapamycin exposure exerted long-lasting protective effects on the intestine by reducing turnover of the epithelium, and preventing age-related increase in ISC proliferation, dysplasia, and loss of intestinal barrier function.

Persisting effects of brief rapamycin treatment could indicate a persistent inhibition of TORC1 activity. S6K is a direct target of TORC1 and reduced phosphorylation of S6K is required for extension of lifespan by rapamycin^2^. Rapamycin treatment instigated later in life, on day 30, reduced TORC1 activity within 48 hours to the same level as chronic treatment in head, muscle, fat body and gut (Extended Data Fig. 2a-d). In contrast to lifespan, terminating rapamycin treatment on day 30 de-repressed TORC1 activity to the level of control flies in all four tissues (Extended Data Fig. 2a-d). In accordance with the absence of a ‘memory effect’ for intestinal S6K phosphorylation, over-expression of constitutively active S6K in the gut did not abolish lifespan extension by chronic or short-term rapamycin treatment (Extended Data Fig. 2e-f and Extended Data Table 4). Thus, TORC1 activity responded acutely to rapamycin, and events downstream of TORC1 other than reduced activity of S6K in the intestine, induced the ‘rapamycin memory’ effects.

Increased autophagy is also a downstream effector of TORC1 and is required for lifespan extension by rapamycin^2^. Persistently up-regulated autophagy could therefore carry the ‘memory of rapamycin’. To assess autophagic flux, we performed co-staining with Cyto-ID and lysotracker dye. While Cyto-ID specifically labels autophagosomes, lysotracker stains autolysosomes, and an increased ratio of autolysosomes to autophagosomes indicates increased autophagic flux ^21^. Chronic rapamycin treatment increased levels of autolysosomes, without altering the levels of autophagosomes, indicative of an increased autophagic flux (Fig 3a). Strikingly, the number of LysoTracker-stained punctae remained fully elevated even 10-days (Fig. 3a) and 30-days (Extended Data Fig. 3) after rapamycin was withdrawn, with no change in Cyto-ID positive punctae (Fig. 3a). Immunoblot analysis revealed that chronic treatment decreased the levels of intestinal, non-lipidated and lipidated forms of the Atg8 protein and the *Drosophila* p62 homolog Ref-2-P and these stayed low 10-days after the treatment from day 1-15 was withdrawn (Fig. 3b), indicative of persistently activated autophagy. However, rapamycin had no effect on Atg8 and Ref-2-P in heads (Fig. 3c), suggesting a tissue-specific response. Together, these results suggest that autophagy induced by brief rapamycin treatment stayed induced for a prolonged period after rapamycin was withdrawn, despite TORC1 activity being restored back to control levels within 48 hours.

**Fig. 3:**
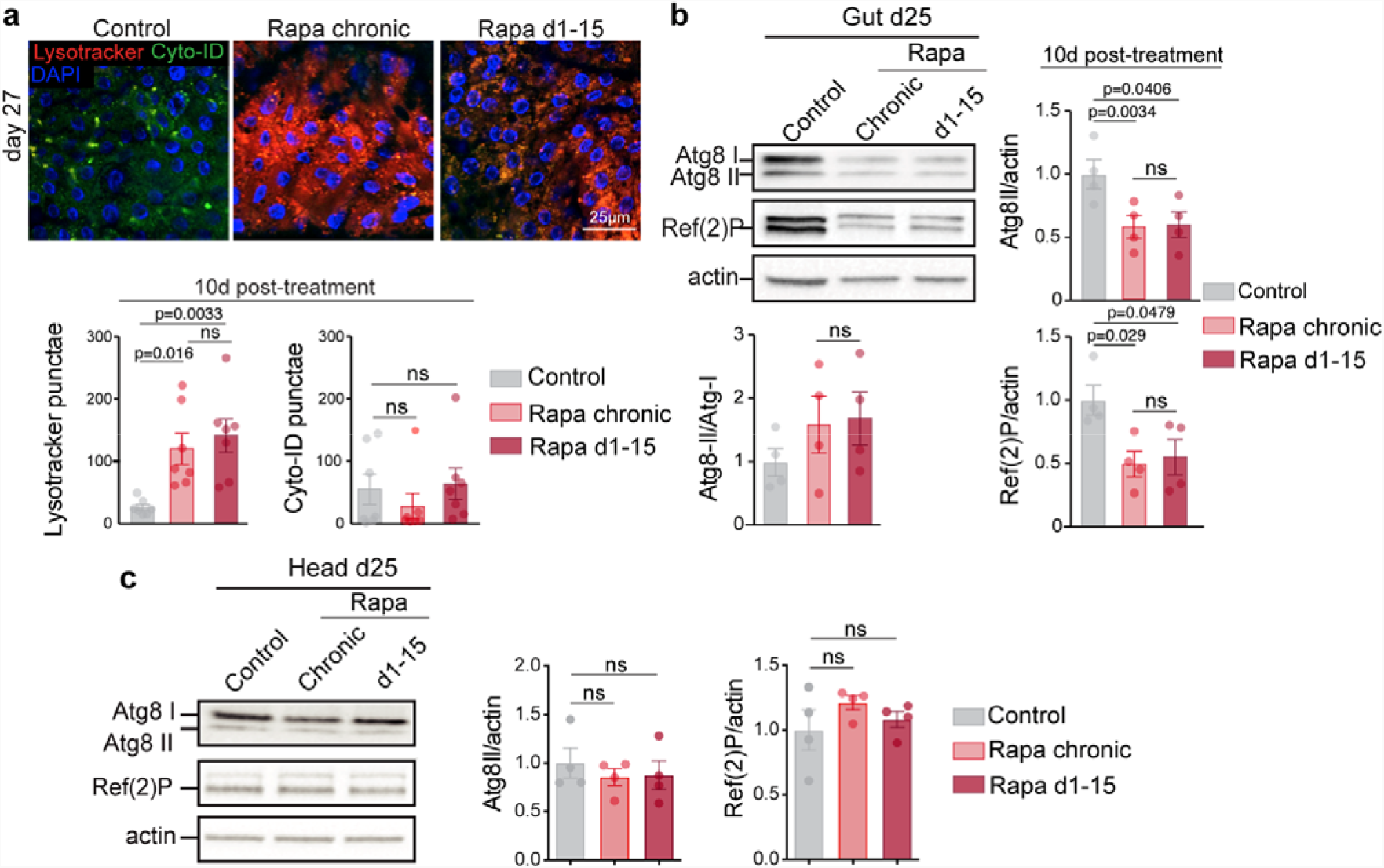
Short-term rapamycin treatment induces lasting autophagy activation. **a**, Number of punctae stained by LysoTracker and Cyto-ID in 25-day old flies chronically or in days 1-15 treated with rapamycin (n=7). **b-c**, Immunoblot of autophagy-related proteins, Atg8-I, Atg8-II and Ref-2-P in the fly gut (**b**) and head (**c**) on day 25, 10-days post-rapamycin-treatment. Data are mean ± s.e.m. One-way ANOVA; Bonferroni’s multiple comparison test. n=4.

To test for a causal role of elevated autophagy in the intestine in the ‘rapamycin memory’, we abrogated it, both briefly and chronically, with double stranded RNA interference. We used inducible GeneSwitch drivers to drive expression of *Atg5*-RNAi in intestinal stem cells (ISCs) or enterocytes. Surprisingly, chronic and day 1-15 treatment with rapamycin both failed to increase lifespan of flies expressing *Atg5*-RNAi specifically in enterocytes (Fig. 4a-b and Extended Data Table 5), but not in ISCs (Fig. 4c-d and Extended Data Table Table 6). Furthermore, enterocyte-specific chronic and day 1-15 over-expression of *Atg5*-RNAi abrogated protection of gut barrier function by chronic and brief rapamycin exposure, respectively (Fig. 4e-f). Blocking the increase in autophagy in response to rapamycin in the enterocytes of the gut thus completely abolished the ‘rapamycin memory’ effect on both lifespan and intestinal integrity.

**Fig. 4.**
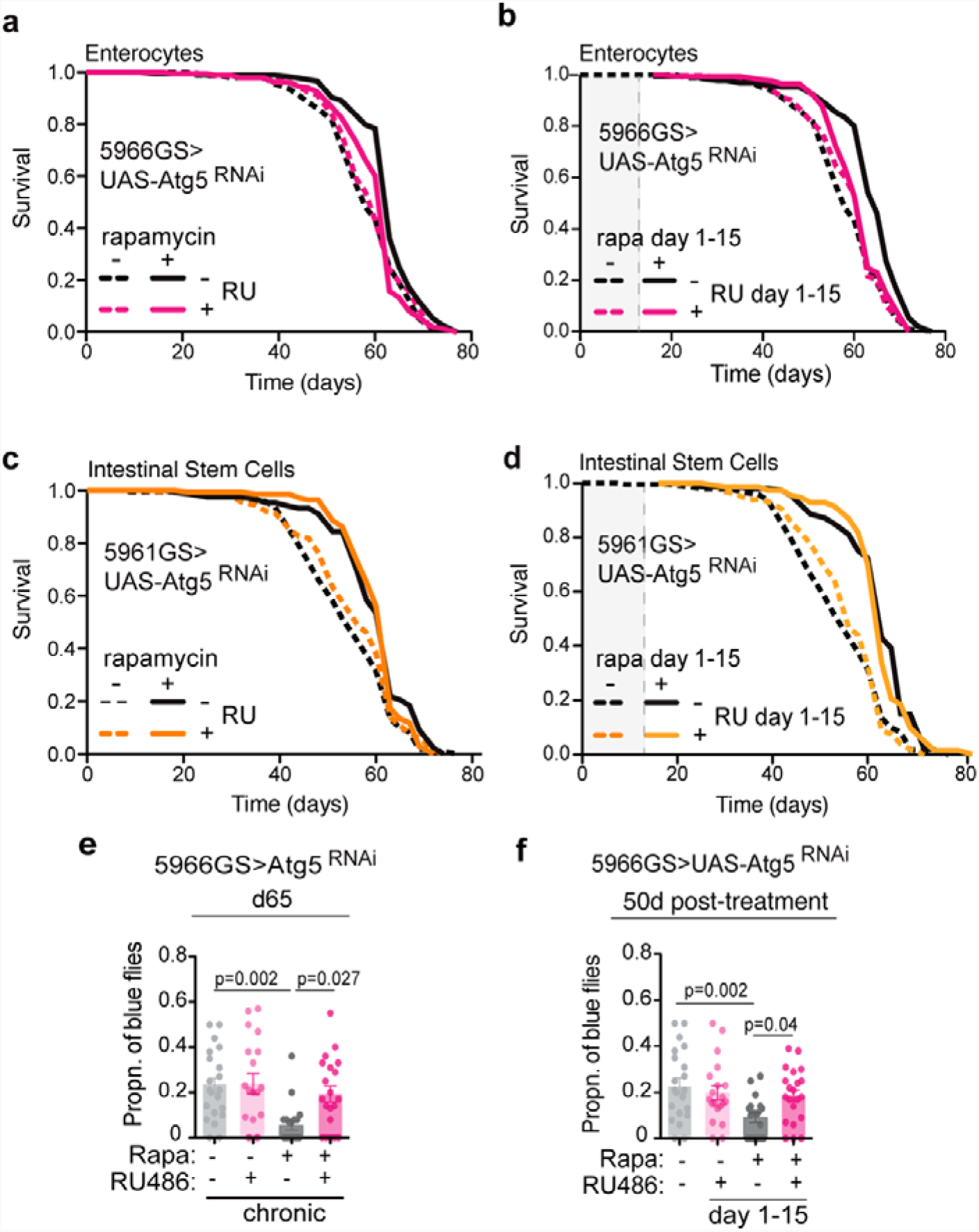
Enterocyte-specific autophagy induction mediates lifespan extension and gut barrier protection by short-term rapamycin treatment. **a-b**, Chronic (p=1.53 × 10^−05^) and brief rapamycin (p=8.8 × 10^−10^) treatment extended lifespan of control flies, but not of flies expressing RNAi against Atg5 in enterocytes (chronic: p=0.25; d1-15: p=0.097, see also Extended Data Table 5). n=400. **c-d**, Chronic (p=3,4 × 10^−06^) and brief rapamycin (p=8.2 × 10^−13^) treatment extended lifespan of control flies and flies with *Atg5*-RNAi specifically in intestinal stem cells (chronic: p =0.001; d1-15: p=5,4 × 10^−12^, see also Extended Data Table 6). n=200. Log-rank test and CPH analysis. **e-f**, Chronic and brief rapamycin treatment reduced the proportion of blue flies in the control group, but not in flies with enterocyte-specific *Atg5*-RNAi. Rapamycin*genotype interaction (chronic: p=0.057; d1-15: p=0.020). n=19-21 vials per condition with 20 flies in each vial. Data are mean ± s.e.m. Two-way ANOVA followed by Bonferroni’s post-test. *p < 0.05, **p < 0.01, ***p < 0.001.

To determine whether direct, genetic activation of autophagy was sufficient to mimic the ‘memory of rapamycin’ in the absence of the drug, we over-expressed *Atg1*, which induces autophagy in flies^22^. Interestingly, similar to rapamycin short-term treatment, over-expression of *Atg1* in enterocytes from days 1-15 caused lasting down-regulation of Ref-2-P 10 days after *Atg1* over-expression was terminated, while combining rapamycin with enterocyte-specific over-expression of *Atg1* from days 1-15 did not further reduce Ref-2-P levels (Fig. 5a). Furthermore, lifelong and day 1-15 enterocyte-specific over-expression of *Atg1* extended lifespan (Fig. 5b-c) and prevented age-related loss of intestinal integrity (Fig. 5d-e) as much as did chronic or brief rapamycin exposure, and the combination of *Atg1* over-expression and rapamycin did not further increase lifespan (Fig. 5b-c and Extended Data Table 7) or improve gut barrier function (Fig. 5d-e). Thus, brief elevation of autophagy in enterocytes induces a memory identical to that from brief rapamycin treatment and mediates the ‘memory of rapamycin’ in increased autophagy, intestinal health, and lifespan.

**Fig. 5:**
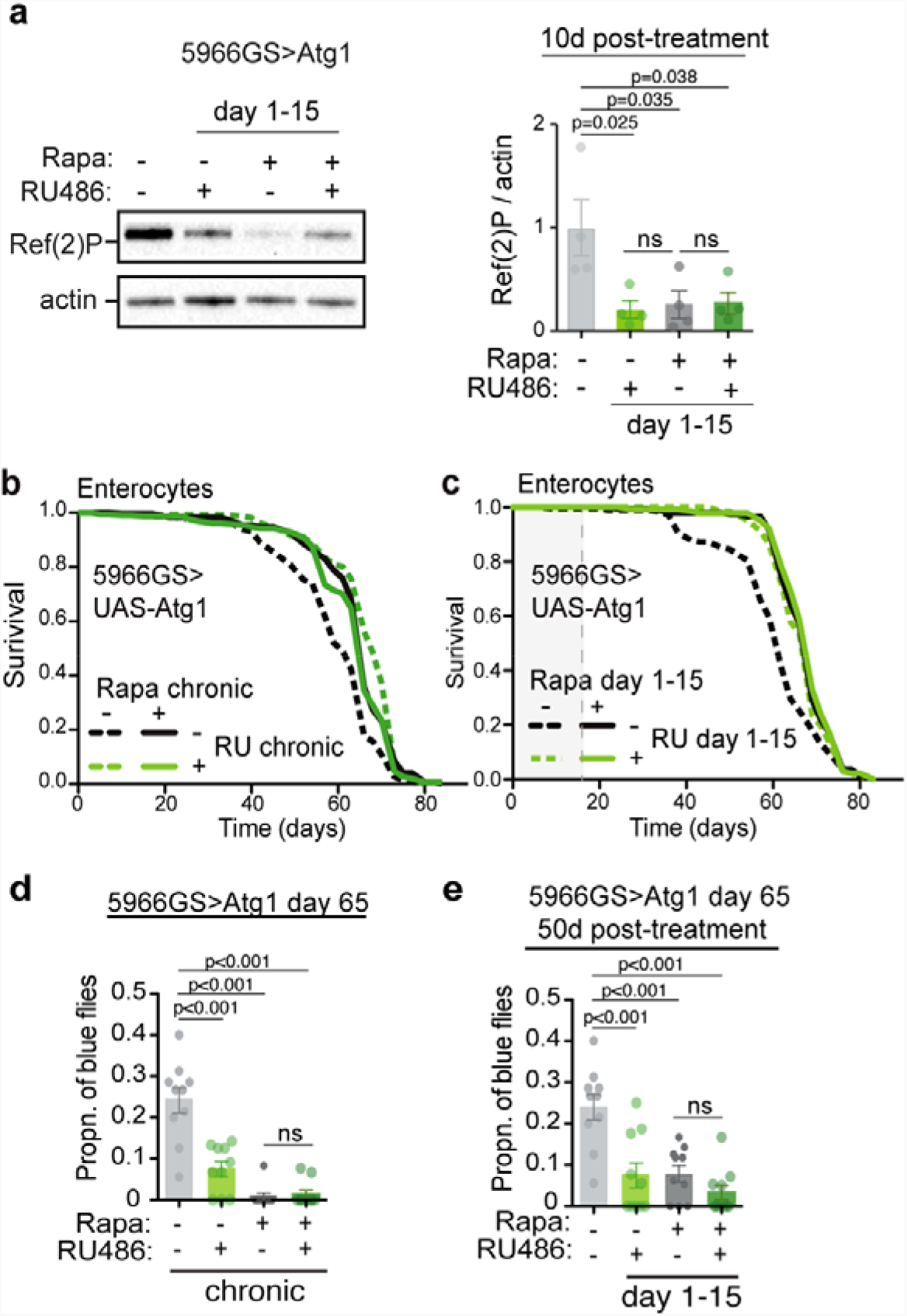
Short-term *Atg1* over-expression induces lasting autophagy activation and extends lifespan to the same degree as short rapamycin treatment. **a**, Immunoblot of intestinal Ref-2-P of flies treated with rapamycin from day 1-15 in combination with enterocyte-specific over-expression of *Atg1* in days 1-15, measured 10 days post-treatment (n=4). Genotype*rapamycin interaction (p=0.03). **b-c**, Chronic (p=3.6 × 10^−11^) and day 1-15 (p=6.7 × 10^−05^) over-expression of *Atg1* specifically in enterocytes extended lifespan to the same degree as rapamycin (Chronic: p=0.50; d1-15: p=0.69, see also Extended Data Table 7). n=160-200. Log rank test and CPH analysis. **g**, Chronic and day1-15 over-expression of *Atg1* reduced the proportion of blue flies to the same degree as rapamycin treatment. Rapamycin*genotype interaction for chronic (p=0.02). n=10 vials per condition with 20 flies per vial. Data are mean ± s.e.m. Two-way ANOVA; Bonferroni’s multiple comparison test.

To test whether the recently reported rapamycin-mediated increase in histone expression ^21^ underlies rapamycin memory, we investigated histone H3 expression after d1-15 rapamycin treatment and the effects of over-expression of H3/H4 during d1-15 on autophagy, gut health and lifespan. As expected, H3 expression and accumulation of chromatin at the nuclear envelope were induced by chronic rapamycin treatment but were decreased back to control levels 15 days post-treatment (Extended data Fig. 4a-b). Moreover, although chronic over-expression of H3/H4 extended lifespan, decreased pH3+ cell count and intestinal dysplasia, and increased lysotracker staining, these phenotypes showed no memory of previous d1-15 H3/H4 expression (Extended data Fig. 4c-f). These data suggest that, although increased histone expression mediates lifespan extension by chronic rapamycin treatment, this mechanism is distinct from the one that is responsible for the memory of short-term rapamycin treatment.

To search for regulators of the ‘memory effect’ of rapamycin and elevated autophagy, we performed proteomics analysis. Gene ontology (GO) term enrichment analysis of the proteins that were increased by rapamycin treatment on day 25 and that remained induced 10 days after the treatment revealed high enrichment in proteins involved in branched-chain amino acid and carbohydrate metabolism, in particular lysosomal mannosidases (Extended Data Fig. 5). We also found an increase in lysosomal alpha-mannosidase V (LManV) mRNA levels by qRTPCR (Extended Data Fig. 6). We therefore tested if knock-down of LManV abolished the ‘memory of rapamycin’. Indeed, knock-down of LManV blocked both the increase in lysotracker-stained punctae by rapamycin treatment during days 1-15 (Fig. 6a) and the improved gut pathology mediated by short-term rapamycin treatment (Fig. 6b). To test if LManV activation was sufficient to mimic short-term rapamycin treatment, we over-expressed LManV during days 1-15, and found that it increased lysotracker-stained punctae and reduced age-related gut pathologies to the same degree as chronic over-expression of LManV (Fig. 6c-d). Taken together these findings suggest that the ‘memory of rapamycin’ in elevated autophagy and improved gut health is mediated through increased expression of LManV.

**Fig. 6:**
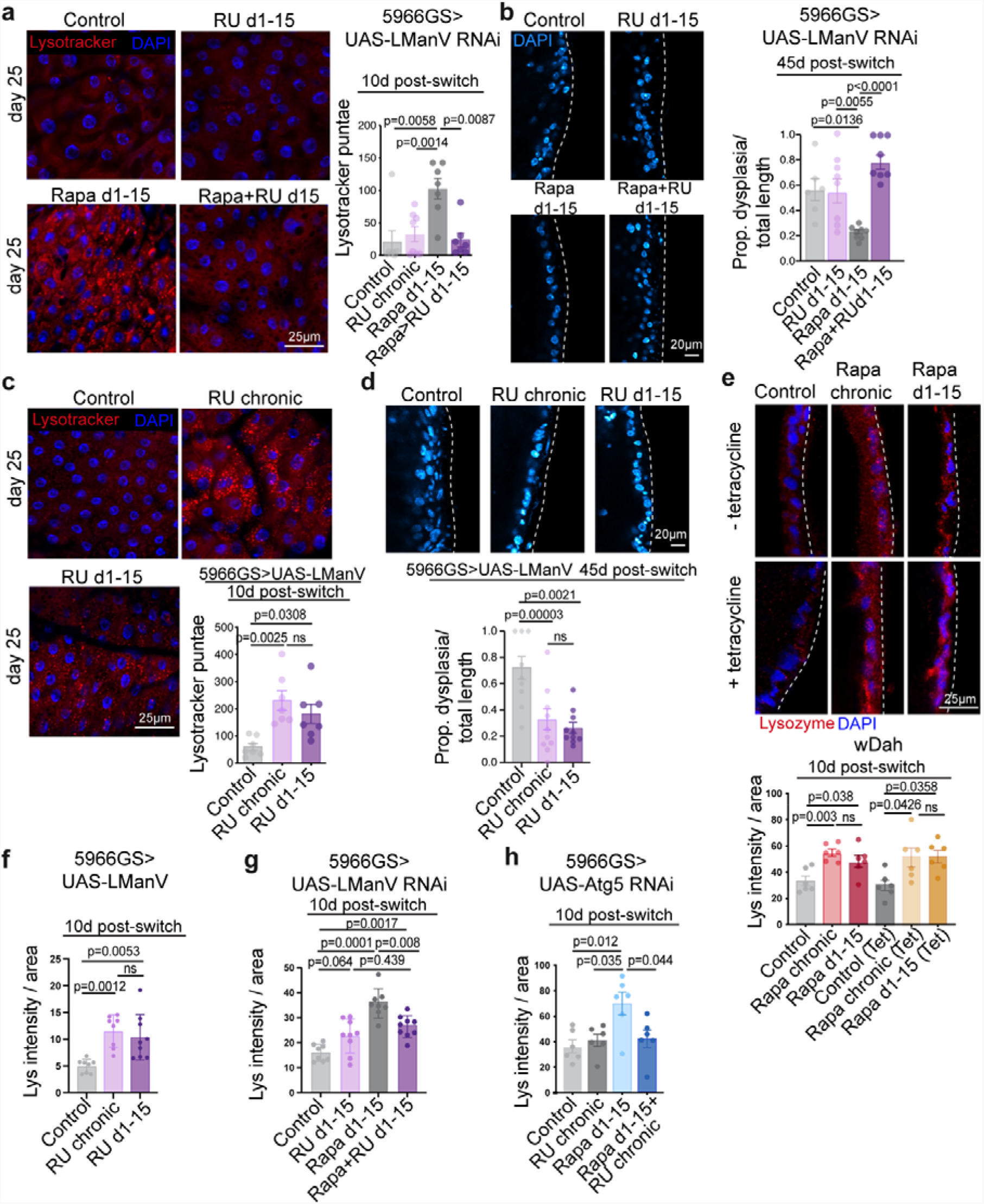
Persistent increase in LManV mediates the “memory of autophagy” and reduced age-related gut pathology induced by short-term rapamycin treatment. **a-b**, Over-expression of RNAi against LManV in enterocytes in days 1-15 abolished increase in lysotracker staining (**a**) and reduction in gut pathology (**b**) induced by short-term rapamycin treatment. **c-d**, Over-expression of LManV in enterocytes in days 1-15 increased lysotracker staining (**c**) and reduced age-related gut-pathology (**d**) to the same degree as chronic over-expression of LManV. **e**, Chronic and short-term rapamycin treatment increased intestinal lysozyme level irrespective of tetracycline treatment. **f**, Chronic and short-term over-expression of LManV increased lysozyme to the same degree. **g**, Over-expression of RNAi against LManV in enterocytes in days 1-15 abolished increase in lysozyme induced by rapamycin treatment in days 1-15. h, Over-expression of RNAi against Atg5 in enterocytes in days 1-15 abolished increase in lysozyme induced by rapamycin treatment in days 1-15. Data are mean ± s.e.m. One-way (**c, d, f**) and Two-way (**a, b, e, g, h**) ANOVA; Bonferroni’s multiple comparison test.

Recent studies showed that lysozyme-associated secretory autophagy plays a key role in gut health and pathogenesis in mammalian small intestine ^23,24^. Secretory autophagy is an autophagy-based alternative secretion system that is activated in response to infection, and it is mediated by core autophagy proteins Atg5 and Atg16L1. Based on our data suggesting the importance of autophagy in the gut for the ‘memory of rapamycin’, we assessed whether levels of intestinal lysozyme, as a proxy for secretory autophagy, were affected by rapamycin treatment. We found that they were increased and remained fully so 10 days after the treatment was withdrawn. These responses to rapamycin were unaffected by tetracycline treatment (Fig. 6e), suggesting that the intestinal microbiota did not play a role. To investigate if LManV and autophagy were responsible for inducing increased lysozyme, we measured lysozyme in intestines of flies over-expressing LManV, and found that both chronic and short-term over-expression increased lysozyme levels to the same degree. Knock-down of LManV by RNAi partially abolished increased lysozyme by rapamycin treatment in days 1-15 (Fig. 6f-g), while blocking autophagy by RNAi against Atg5 abolished the increase in lysozyme induced by d1-15 rapamycin treatment (Fig 6h). Together, these data suggest that autophagy and LManV mediate rapamycin-induced increase in intestinal lysozyme.

Branched-chain amino acid aminotransferase (BCAT) is one of the enzymes catabolizing the first step of BCAA degradation and we therefore tested if knock-down of BCAT also abolished the ‘memory of rapamycin’. Expression of RNAi against BCAT in enterocytes from day 15 onwards blocked the increased number of lysotracker-stained punctae and partially blocked improved gut pathology and lifespan extension achieved by short rapamycin treatment (Extended Data Fig. 7a-c). Taken together, these findings suggest that BCAT contributes to the ‘memory of autophagy’ and partially mediates the effects of rapamycin on gut health and longevity.

To assess if lasting benefits of a short-term rapamycin treatment are conserved between flies and mammals, we assessed the impact of rapamycin treatment on intestinal permeability in mice (Fig. 7a), by measuring plasma lipopolysaccharide-binding protein (LBP) levels, a marker of bacterial translocation from intestine into circulation^25,26^. As we (Fig. 7b) and others^27^ showed that the age-related increase in gut permeability in rodents appears already in middle-age, we treated mice with rapamycin chronically or from 3-6 months of age, and collected samples 6 months after the treatment was withdrawn, at 12-months of age (Fig. 7a). Strikingly, 6 months after rapamycin was withdrawn plasma LBP levels were reduced to levels similar to those with chronic treatment, (Fig. 7b), suggesting that the long-lasting, beneficial effects of short-term rapamycin exposure on intestinal integrity is conserved in mammals.

**Fig. 7:**
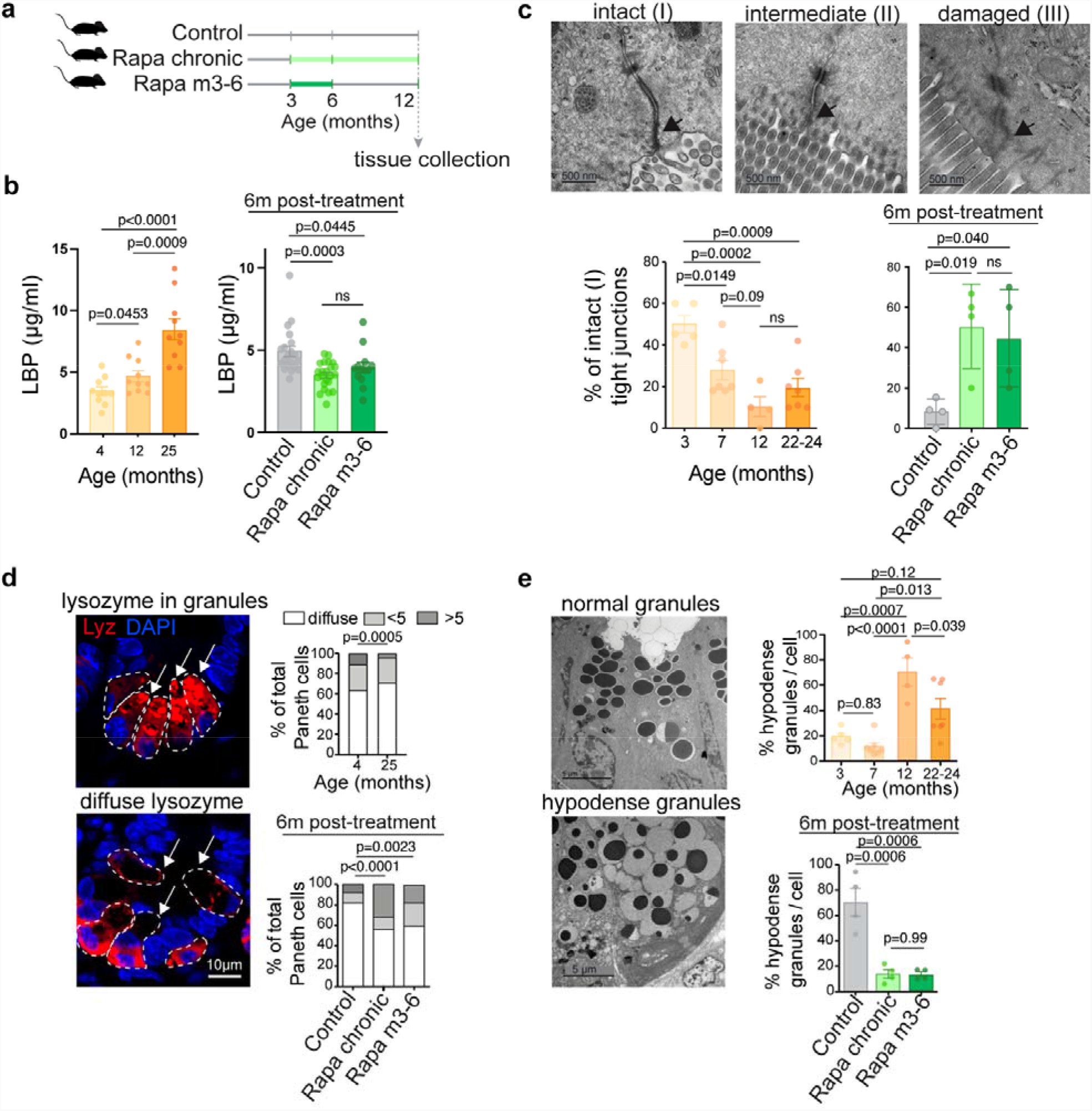
Short-term rapamycin exposure maintains gut barrier function and Paneth cell architecture to the same degree as lifelong treatment in mice. **a**, Experimental design. **b**, Plasma LBP levels during ageing and 6 months after rapamycin treatment was terminated (n=10-15 mice). **c**, Tight junction pathology score: I - narrow and electron dense TJs; II - reduced electron density, but no dilations within TJs; III - low electron density and dilated TJs. Proportion of intact TJs during ageing and 6 months post-rapamycin-treatment. **d**, The proportion of Paneth cells with diffuse lysozyme staining was increased in aged mice and rapamycin reduced the proportion of Paneth cells (arrows) with diffuse lysozyme staining, which remained reduced 6 months post-treatment (n=4). **e**, Proportion of hypodense Paneth cell granules in mouse jejunum during ageing and 6 months after rapamycin treatment was withdrawn. Data are mean ± s.e.m (b, e). and s.d (c, rapa treatment). One-way ANOVA; Bonferroni’s multiple comparison test.

Increased gut permeability is associated with compromised tight junctions (TJ)^28^. Irregularities of TJ can be observed by electron microscopy as reduced electron density of the perijunctional ring^29^ and dilations within tight junctions^30^. We analyzed ultrastructure of TJs in jejunal villi. Intact TJs, which appeared as narrow and electron-dense structures, were classified as class I, narrow TJs with reduced electron density, but without dilations within the TJ as class II, and TJs that were both low in electron density and dilated as class III (Fig. 7c). In line with previously published data on gut permeability ^27^, TJ quality declined during ageing, with 7 month old mice already showing reduced proportion of intact TJs compared to 3 month old mice (Fig. 7c). In accordance with plasma LBP result, rapamycin treatment increased the proportion of intact TJs, which remained increased 6 months after rapamycin withdrawal, further supporting the hypothesis that rapamycin protects age-related decline in intestinal integrity (Fig. 7c and Extended Data Fig. 8).

Paneth cells are specialized secretory cells that serve as a niche for ISCs^31^ and contain secretory granules filled with antimicrobial proteins, such as lysozyme, and rapamycin improves the Paneth cell function and their support of ISCs^32^. Lysozyme is normally efficiently packed in Paneth cell granules^23^. In 12 months-old control mice, we observed a notable proportion of Paneth cells with abnormal lysozyme distribution, which was diffuse in cells. Short-term rapamycin treatment increased the proportion of cells with lysozyme packed granules and reduced those with a diffuse lysozyme signal (Fig. 7d). Transmission electron microscopy further showed that Paneth cell granule abnormalities, seen as loosely packed and hypodense granules that are a feature of dysfunction^23^, appeared already at 12 months of age (Fig. 7e). Remarkably, rapamycin treatment decreased the proportion of hypodense Paneth cell granules, which stayed decreased to the levels seen with chronic treatment 6 months after rapamycin treatment was withdrawn (Fig. 7e). Together, these data suggest that short-term rapamycin treatment abolished age-related Paneth cell abnormalities^23^.

Next, we assessed if the long-term elevation of autophagy by past rapamycin treatment is conserved in mice. Although chronic and 3-6 months rapamycin treatment did not significantly reduce the number of p62 punctae in the villi region, comprising enterocytes and goblet cells, there was a trend in the villi of 12 months old rapamycin treated mice (Extended Data Fig. 9). As autophagy is essential for proper Paneth cell function and secretion^23^, and upon autophagy activation autophagy-related proteins colocalize with Paneth cell granules^33^, we measured the number of granules positive for both lysozyme and p62. Chronic and 3-6 months rapamycin treatment increased the number of Paneth cell granules positive for both lysozyme and p62 assessed at 12 months of age (Fig. 8a), suggesting that autophagy in Paneth cells may play a key role in improving cell health in response to brief treatment, even 6 months after the drug withdrawn.

**Fig. 8:**
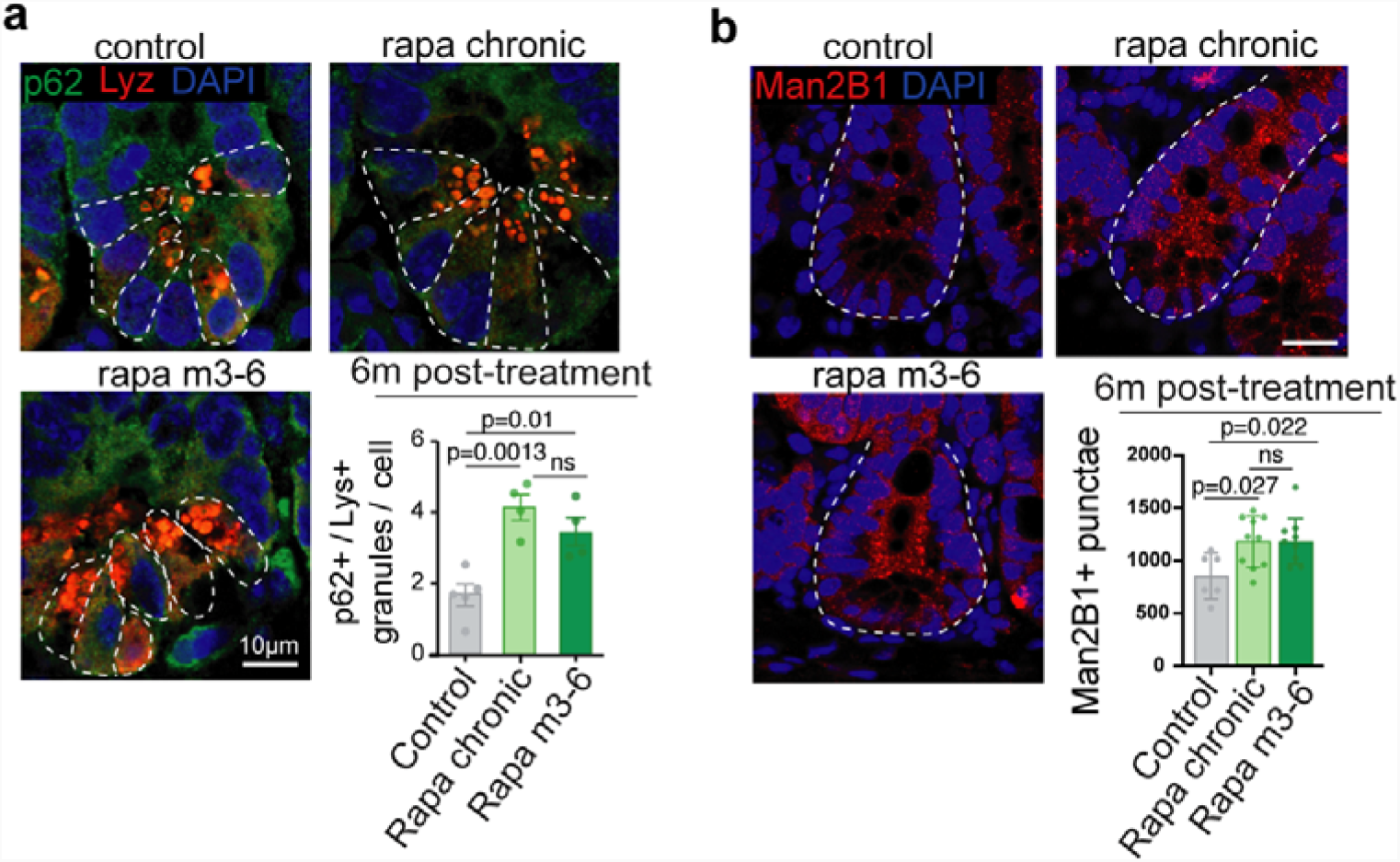
Short-term rapamycin exposure increases the number of granules positive for both lysozyme and p62 in Paneth cells and the number of Man2B1 positive punctae in intestinal crypts to the same degree as lifelong treatment in mice. **a**, The number of lysozyme+/p62+ granules per Paneth cell was increased by rapamycin and remained increased 6 months after the treatment was withdrawn (n=4). White dashed denotes a Paneth cell. **b**, Number of Man2B1+ punctae was increased by rapamycin and remained increased 6-months post-treatment. White dashed denotes a crypt unit. Data are mean ± s.e.m. One-way ANOVA; Bonferroni’s multiple comparison test.

As we showed that LManV is one of the mediators of rapamycin memory in *Drosophila*, and that it also mediates rapamycin-induced increase in lysozyme levels in flies, we measured the levels of mannosidases in mouse gut. We observed that rapamycin increased the number of Man2B1 positive punctae in intestinal crypts, and these stayed increased 6 months after the treatment was withdrawn and to the same degree as with chronic treatment (Fig 8b), in line with the fly data.

Paneth cell health is critical to the homeostasis of the small intestine, including promoting intestinal stem cell proliferation and maintenance, which eventually mediates regenerative capacity of the intestinal epithelium^34,35^. We measured the regenerative ability by assessing mouse intestinal epithelial crypts to form clonogenic organoids *in vitro*. We treated mice with rapamycin starting from 15-21 months of age, followed by a switch to control food for another 2 months (Extended Data Fig. 10a). Interestingly, short-term treatment increased organoid forming potential of intestinal crypts isolated 2 months after drug withdrawal (Extended Data Fig. 10b). Regenerative growth of de novo crypts was also increased in organoids generated from intestines from pre-treated mice (Extended Data Fig. 10c).

Together, these data show that short-term rapamycin exposure in adult mice combated age-related decline in intestinal TJ structure, Paneth cell architecture and gut barrier function, and that these geroprotective effects were equivalent to those seen with chronic drug exposure and lasted long after rapamycin treatment was withdrawn. In addition, our data indicate that brief rapamycin may improve regenerative capacity of the intestinal epithelium long-term.

Our study has uncovered a long-lasting effect of short-term rapamycin administration, including prolonged activation of autophagy, reduced age-related gut pathologies and extension of lifespan in *Drosophila*. Brief rapamycin administration in early adult life induced these benefits to the same degree as lifelong treatment, with a key role of the enterocytes in the intestine. The long-term elevation in autophagy was mediated by the lasting increase in LManV and BCAT expression. Importantly, some of these benefits from early, brief rapamycin treatment were also observed in the small intestine of mice, suggesting the ‘rapamycin memory’ is at least partially conserved in this mammalian model. These findings are intriguing in light of the key role of autophagy in an array of age-related diseases, including cancer^36^, immune system dysfunction^8^, and neurodegenerative diseases^37^. Our findings suggest that the geroprotective effects of rapamycin can be achieved by early, short-term treatment, without the adverse effects sometimes seen with chronic, long-term dosing. While our data shed light on a new path to achieve geroprotection via pharmacological interventions, it will be important to determine the temporal clinical dosing regimen that maximizes protection while minimizing side-effects.

## Methods

### Fly husbandry and strains

The white Dahomey (w^*Dah*^), Wolbachia positive strain was used, unless otherwise stated. Fly stocks were maintained at 25°C on a 12 h light/dark cycle, at constant humidity (60%), and reared on sugar/yeast/agar (SYA) diet, at standard larval density, by collecting eggs on grape juice plates, washing with PBS and pipetting 20 μl of the eggs into each culture bottle. Eclosing adult flies were collected over 18 h and mated for 48 hours, then sorted into single sexes. Female flies were used. All mutants and transgenes were backcrossed for at least six generations into the w^*Dah*^ background, except UAS-BCAT-RNAi line. The following strains were used in the study: *TiGS* ^38^, *5966GS* ^39^, *5961GS* ^15,40^, *Resille-GFP* from the Flytrap project^41^, *UAS*-*Atg5*-RNAi and *UAS*-*Atg1* OE (GS10797) obtained from the Kyoto Drosophila Genetic Resource Center ^42,43^, *UAS*-*LManV* ^44^, *UAS*-*LManV RNAi* (GD13040) obtained from Vienna *Drosophila* Stock Center, UAS-BCAT RNAi (38363) obtained from Bloomington Drosophila Stock Center, *UAS-H3/H4* generated in this lab ^21^.

### Standard media and rapamycin treatment for *Drosophila*

Standard SYA medium was used, containing per liter (L) 100 g autolyzed yeast powder (brewer’s yeast, MP Biomedicals), 50 g sucrose (Sigma), 15 g agar (Sigma), 3 ml propionic acid (Sigma), 30 ml Nipagin (methyl 4-hydroxybenzoate) and distilled water to 1 L. SYA diet was prepared as described before^45^. Rapamycin was dissolved in ethanol, and added to the food in concentration of 200 μM.

### Lifespan assays

Females were placed into vials containing experimental diets and drugs, at a density of 20 flies/vial, and transferred into vials containing fresh food every 2-3 days, when the number of dead flies was scored. Sample size and analyses of all lifespan data are shown in Extended Data Tables 1-7.

### Mouse husbandry and rapamycin treatment

Female C3B6F1 hybrids were used and were bred in an in-house animal facility at the Max Planck Institute for Biology of Ageing. C3B6F1 hybrids were generated by a cross between C3H female and C57BL/6J male mice, obtained from Charles River Laboratories. Four-week-old mice were housed in individually ventilated cages, in groups of five mice per cage, under specific-pathogen-free conditions. Mice had *ad libitum* access to chow (Ssniff Spezialdiäten GmbH; 9% fat, 24% protein, 67% carbohydrates) and drinking water at all times. Experiments were conducted under the approval of the State Office for Nature, Environment and Consumer Protection North Rhine-Westphalia (approval nos. 84-02.04.2017.A074 and 84-02.04.2015.A437). For 6-months post-switch measurements, rapamycin was added at concentration of 14 ppm (mg of drug per kg of food), encapsulated in Eudragit S100 (Evonik). Control chow contained Eudragit encapsulation medium only. Rapamycin treatment was initiated at 3 months of age and was administered either continuously until 12-months of age (rapamycin chronic group) or until month 6, after which the switch-off group received control chow for an additional 6 months (rapamycin 3-6M group). All mice from 6-months-post-switch-experiment were sacrificed at 12 months of age. For the 2-month post-switch organoid experiment, rapamycin treatment (42 ppm, -week-on/1-week-off intervals) was started at 15 months of age and terminated at 21 moths of age, after which switch-off group received control chow for an additional 2 months. Mice were sacrificed at 23 months of age and 2 months post-treatment.

### Western blot analysis

Tissues were lysed in 2xLaemmli buffer (head, thorax and fat body) and proteins denatured at 95°C for 5 min. Proteins from gut were extracted using in 20% trichloric acid, washed in 1 M Tris buffer (not pH’d), resuspended in 2xLaemmli buffer and denatured at 95°C for 5 min. Proteins (10 μg) were separated using pre-stained SDS-PAGE gels (Bio-Rad) and wet-transferred onto 0.45 μm nitrocellulose membrane (GE Healthcare). Blots were incubated with primary p-T389-S6K (CST, 9209), S6K^2^, Atg8 and Ref-2-P^46^ antibodies. Appropriate HRP-coupled secondary antibodies (ThermoFisher) were used. Signal was developed using ECL Select Western Blotting Detection Reagent (GE Healthcare). Images were captured using ChemiDocImager (Bio-Rad) and band intensity was analyzed using ImageJ.

### Immunostaining of fly intestines

Flies were immobilized on ice and guts were dissected in ice-cold PBS. Dissected guts were immediately fixed in 4% formaldehyde for 30 min, washed in 0.2% Triton-X / PBS (PBST) and blocked in 5% bovine serum albumin (BSA) / PBS for 1 h on a shaker. Gut tissues were incubated with primary pH3 (CST, 9701; 1:500), dpErk (CST, 4370;1:400) or lysozyme (ThermoFisher Scientific, PA5-16668; 1:100) solutions in 5% BSA overnight at 4°C, followed by incubation in secondary Alexa Fluor 594⍰donkey anti-rabbit antibody (ThermoFisher Scientific, A2s207; 1:1000). Guts were mounted in mounting medium containing DAPI (Vectashield, H1200), scored and imaged using a Leica inverted microscope for the cell division assay and confocal SP8-DLS for the dpErk staining. pH3 and dpErk imaging was done on the R2 region proximal to proventriculus and for each intestine 3 adjacent images were taken.

### Gut turnover assay

*w*^*Dah*^ were crossed to the *esg*^*ts*^ F/O flies (w; *esg*-*Gal4, tubGal80*^*ts*^, *UAS*-*GFP*; *UAS*-*flp, Act>CD2>Gal4*). Crosses were maintained and progeny were raised at 18°C. Following a 3-day mating at 18°C, female flies were distributed into vials containing EtOH or rapamycin and kept at 18°C for 15 days. On day 15, a subgroup of flies was switched from rapamycin to EtOH food and all experimental groups were transferred to 29°C. Flies were maintained at 29°C for 10 and 20 days, after which guts were dissected, fixed in 4% formaldehyde, and mounted in DAPI-containing mounting medium (Vectashield, H1200). Samples were imaged under a confocal microscope (Leica TCS SP8-X), and images analyzed using ImageJ. The GFP-marked regions represent ISCs and their newly generated progenitor cells, and the GFP-marked area compared to the total corresponding gut area indicates the gut turnover rate. Images were obtained from R4 and R5 intestinal regions.

### Gut barrier analysis

Flies were aged for 65 days on standard SYA diet then transferred into vials containing SYA food with 2.5 % (w/v) FD&C blue dye no. 1 (Fastcolors). The proportion of blue (whole body is blue) or partially blue (at least 2/3 of body is blue) flies was scored 24 h after exposure to the blue food.

### Imaging of gut dysplasia

Guts were dissected in ice-cold PBS, fixed in 4% formaldehyde for 30 min and mounted in DAPI-containing mounting medium (Vectashield, H1200). Endogenous GFP and DAPI were imaged using a confocal microscope. For each condition, 6-14 guts were imaged. The area affected by tumors was measured using the measure function in ImageJ, and the average proportion of the affected area for each gut was calculated.

### Lysotracker staining, imaging and image analysis

Flies were immobilized on ice, dissected in PBS and stained with 0.5 μM Lysotracker DS Red DND-99 (Invitrogen, Molecular Probes), containing 1:2000 0.5 mg/ml Hoechst 33342 (Sigma) for 3 min in 12-well plates on a shaker. Immediately after staining, guts were mounted (Vectashield, H1000) and imaged using a Leica SP8-X confocal microscope. For each gut preparation, an area proximal to the proventriculus was imaged to control for variation across different gut regions, and 3 adjacent images per gut were captured. Images were analyzed using IMARIS software. This experiment was carried out under blinded conditions.

### LBP measurement in mouse plasma

Lipopolysaccharide-binding protein (LBP) was measured in mouse plasma samples by ELISA according to the manufacturer’s instructions (HyCult Biotech, HK: 205).

### Transmission electron microscopy

The intestine was fixed in 2 % glutaraldehyde / 2 % formaldehyde in 0.1 M cacodylate buffer (pH 7.3) for 48 h at 4°C. Afterwards, samples were rinsed in 0.1 M cacodylate buffer (Applichem) and post-fixed with 2 % osmiumtetroxid (Science Services) in 0.1 M cacodylate buffer for 2 h at 4°C. Samples were dehydrated through an ascending ethanol series (Applichem) and embedded in epoxy resin (Sigma-Aldrich). Ultrathin sections (70 nm) were cut with a diamond knife (Diatome) on an ultramicrotome (EM-UC6, Leica Microsystems) and placed on copper grids (Science Services, 100mesh). The sections were contrasted with 1.5 % uranyl acetate (Plano) and lead citrate (Sigma-Aldrich). Images were acquired with a transmission electron microscope (JEM 2100 Plus, JEOL) and a OneView 4K camera (Gatan) with DigitalMicrograph software at 80 KV at RT. For each mouse and for each measured phenotype, 10 random images were taken and the final score for each mouse was calculated as a mean value obtained from 10 images. Imaging and scoring of EM data were carried out under blinded conditions.

### Isolation of mouse intestinal crypts and organoid culture

Mouse jejunal section were used to isolate crypts according to the manufacturer’s instructions (STEMCELL Technologies, document 28223). Complete IntestiCult medium was exchanged every 2-3 days and organoid numbers and *de novo* crypts were scored on days 5 and 7. This experiment was carried out under blinded conditions.

### Immunostaining of mouse tissues

Jejunal sections were fixed in 4 % PFA, embedded in paraffin and sectioned. Slides were deparaffinized and antigen retrieval was performed by boiling with pH 6 citrate buffer. Primary antibodies used were: p62/SQSTM (Abcam, 56416; 1:100), pH3 (CST, 4370; 1:100), lysozyme (ThermoFisher Scientific, PA5-16668; 1:300), Man2B1 (St John’s Laboratory, 640-850; 1:100). Primary antibodies were detected using Alexa Flour 488-, Alexa Flour 594-and Alexa Flour 633-conjugated anti-rabbit or anti-mouse secondary antibodies (Thermo Fisher Scientific, 1:500). Sections were mounted in DAPI-containing mounting medium (Vectashield H-1200) and imaged using confocal Leica SP8-DLS or SP8-X microscope.

### Metabolite extraction for lipid and polar metabolites from a single sample

A fixed volume of 750 µL of a mixture of methanol: acetonitrile: water (40:40:20 [v:v]), containing the appropriate internal standard (Everolimus, LC Laboratories) was added to the samples. Samples were incubated for 30 min on an orbital shaker at 4°C followed by sonication for 10 min in a cooled bath-type sonicator. Subsequently, samples were centrifuged for 10 min at 16.000x g at 4°C and the supernatants were collected, while the insoluble pellet was dried by aspirating the remaining liquid. 600 µL of chloroform: water (2:1 [v:v]) was added to the collected supernatants and samples were mixed on the orbital shaker at 4°C for 10 min. Phases were separated by centrifugation at 16.000x g and 4°C for 5 min. 500 µL of the upper, polar phase and 500 µL of the lower, hydrophobic chloroform phase were collected and concentrated in a speed vac concentrator.

### Rapamycin analysis by targeted UPLC-MS

The concentrated chloroform-phase was resuspended in 50 µL of an acetonitrile:isopropanol:DMSO [70:30:5 [v:v]) solution and the samples were sonicated for 5 min in a cooled bath-type sonicator. Insoluble components were removed by 10 min centrifugation at 16000x g at 4°C and cleared supernatant was transferred to a fresh auto sampler tube and used for the UPLC (Acquity i-class, Waters) -MS (TQ-S, Waters) analysis. Chromatographic separation of 2 µL of the extract was performed using a 2.1 mm x 100 mm BEH C8 column (Waters), using buffer A containing 10 mM ammonium acetate and 0.1% acetic acid in water. Buffer B consisted of a mixture of 70:30 acetonitrile:isopropanol [v:v] containing 10 mM ammonium acetate and 0.1 % acetic acid. The gradient, which was carried out at a flow rate of 300 µL/min was 0 min 55% A, 0.5 min 55 % A, 1.5 min 0 % A, 3.5 min 0 % A, 3.6 min 55 % A and 6 min 55 % A. Samples were analysed by MRM analysis, monitoring the main adducts (NH4) of Everolimus and Rapamycin. The selected transitions for each compound were m/z 931.78 ➔ 864.64 and 931.78 ➔ 882.64 for Rapamycin, while Everolimus was monitored using transitions 975.65 ➔ 908.5 and 975.65 ➔ 926.5. Data analysis was performed using TargetLynx XS.

### Peptide generation and TMT-labeling

20 μL of lysis buffer (6 M GuCl, 2.5 mM TCEP,10 mM CAA, 100 mM Tris-HCl) was added to 25 guts and tissues were homogenized using a hand-homogenizer. Homogenates were heated at 95°C for 10 min and subsequently sonicated using the Bioruptor (10 cycles, 30 sec sonication/30 sec break, high performance). Samples were centrifuged for 20 min at 2000x g and supernatant was diluted 10-fold in 20 mM Tris. Protein concentration in the supernatant was measured using a NanoDrop and 1:200 (w/w) of trypsin (Promega, Mass Spectrometry grade) was added to 200 µg of sample. Trypsin digestion was performed overnight at 37°C and stopped by the addition of 50% of formic acid (FA) to a final concentration of 1 %. Peptide clean-up was carried out using an OASIS HLB Plate. Wetting of the wells was performed by the addition of 200 µl of 60 % acetonitrile/0.1 % FA and equilibration adding 400 µl of 0.1 % FA. The sample and 100 µl of 0.1 % FA were loaded into the wells and peptides eluted by the addition of 80 µl of 60 % ACN/0.1 % FA. Peptides were air-dried by using the SpeedVac and the pellet resuspended in 60 µl of 0.1% FA. 15 µg of peptides was dried in SpeedVac and used for tandem mass tag (TMT) labelling. The pellet was dissolved in 17 µl of 100 mM triethylammonium bicarbonate (TEAB) and 41 µl of anhydrous acetonitrile was added. Samples were incubated for 10 min at RT with occasional vortexing, followed by the addition of 8 µl of TMT label and subsequent incubation for 1 h at RT. The labelling reaction was stopped by the addition of 8 µl of 5 % hydroxylamine and incubation for 15 min. Samples were air-dried in SpeedVac, resuspended in 50 µl of 0.1 % FA and cleaned with an OASIS HLB Plate as previously described. 4 replicates per condition and 25 intestines per replicate were used for peptide generation and TMT-labelling for proteomics analysis.

### High-pH fractionation

Pooled TMT labeled peptides were separated on a 150 mm, 300 μm OD, 2 μm C18, Acclaim PepMap (Thermo Fisher Scientific) column using an Ultimate 3000 (Thermo Fisher Scientific). The column was maintained at 30°C. Buffer A was 5 % acetonitrile 0.01M ammonium bicarbonate, buffer B was 80 % acetonitrile 0.01M ammonium bicarbonate. Separation was performed using a segmented gradient from 1 % to 50 % buffer B, for 85min and 50 % to 95 % for 20 min with a flow of 4 μL. Fractions were collected every 150 sec and combined into nine fractions by pooling every ninth fraction. Pooled fractions were dried in a Concentrator plus (Eppendorf), resuspended in 5 μL 0.1% formic acid from which 2 μL was analyzed by LC-MS/MS.

### LC-MS/MS analysis

Peptides from each of the nine high-pH fractions were separated on a 25 cm, 75 μm internal diameter PicoFrit analytical column (New Objective) packed with 1.9 μm ReproSil-Pur 120 C18-AQ media (Dr. Maisch,) using an EASY-nLC 1200 (Thermo Fisher Scientific). The column was maintained at 50°C. Buffer A and B were 0.1 % formic acid in water and 0.1 % formic acid in 80 % acetonitrile. Peptides were separated on a segmented gradient from 6% to 31% buffer B for 120 min and from 31v% to 50 % buffer B for 10 min at 200 nl/min. Eluting peptides were analyzed on an Orbitrap Fusion mass spectrometer (Thermo Fisher Scientific) in TMT-SPS mode. Peptide precursor m/z measurements were carried out at 60000 resolution in the 350 to 1500 m/z range with an AGC target of 1e6. Precursors with charge state from 2 to 7 only were selected for CID fragmentation using 35 % collision energy and an isolation window width of 0.7. The m/z values of the peptide fragments, MS/MS, were measured in the IonTrap at a “Rapid” scan rate, a minimum AGC target of 1e4 and 100 ms maximum injection time. Upon fragmentation, precursors were put on a dynamic exclusion list for 45 sec. The top ten most intense MS/MS peaks were subjected to multi-notch isolation with an AGC target of 5e4 and 86 ms maximum injection time and further fragmented using HCD with 65 % collision energy. The m/z values of the fragments, MS3, were measured in the Orbitrap at 50 K resolution. The cycle time was set to two seconds.

### Protein identification and quantification

The raw data were analyzed with MaxQuant version 1.5.2.8^47^ using the integrated Andromeda search engine^48^. Peptide fragmentation spectra were searched against the canonical and isoform sequences of the *Drosophila melanogaster* reference proteome (proteome ID UP000000803, downloaded September 2018 from UniProt). Methionine oxidation and protein N-terminal acetylation were set as variable modifications; cysteine carbamidomethylation was set as fixed modification. The digestion parameters were set to “specific” and “Trypsin/P,” The minimum number of peptides and razor peptides for protein identification was 1; the minimum number of unique peptides was 0. Protein identification was performed at peptide spectrum matches and protein false discovery rate of 0.01. The “second peptide” option was on. The quantification type was set to “Reporter ion MS3” and “10-plex TMT”. Prior to the analysis, the TMT correction factors were updated based on the values provided by the manufacturer.

### Bioinformatics

#### Proteomics data analysis

Intensity values were log2 transformed and each sample was separately z-transformed. For simpler interpretation the z-scores were rescaled to approximately their original scale by multiplying each z-score with the overall standard deviation of the original log2 transformed data and adding back the overall mean of the original log2 transformed data. The normalized data were filtered for proteins that were detected in at least three replicates per biological group and proteins annotated as contaminant or reverse identification were removed. Missing values after filtering were imputed using the impute.knn function from the impute package version 1.56.0^49^. Differential expression analysis was performed using the limma package version 3.38.3^50^. P-values were corrected for multiple testing using the Benjamini-Hochberg procedure and a significance threshold of 0.05 was used to determine significant differential expression. Differential expression was determined between the following biological groups: 25-day old flies chronically treated with rapamycin vs. 25-day old control flies and 25-day old flies treated with rapamycin from day 1-15 vs. 25-day old control flies. The normalized data after batch effect removal with the removeBatchEffect function from the limma package was used for principal component analysis using the prcomp function from the stats R package version 3.5.3.

### Gene ontology term enrichment

The topGO package version 2.32.0****^51^ with the annotation package org.dm.e.g.db^52^ was used for Gene ontology term enrichment analysis. The weight01 Fisher procedure^53^ was used with a minimal node size of five. The enrichment of each term was defined as the log2 of the number of significant genes divided by the number of expected genes. Protein groups of interest were tested for enrichment against a universe of all detected proteins. Only significantly enriched terms with a minimum of three significant proteins and a maximum of 300 annotated genes were used in the cell plot

### Statistical analyses

Statistical analysis was performed in GraphPad Prism except for survival analysis. Statistical tests for each experiment are mentioned in the corresponding figure legends. Survival data were analyzed with Log-rank test and Cox Proportional Hazard analysis, using Excel (Microsoft) and Jmp (SAS Institute) software, respectively. Bioinformatics analysis was performed using R (version 3.5.5).

## Data Availability

The mass spectrometry proteomics data have been deposited to the ProteomeXchange Consortium via the PRIDE partner repository with the dataset identifier PXD020820. Source data are presented with the paper.

## Acknowledgments

We thank the core facilities at the Max Planck Institute for Biology of Ageing: Christian Kukat and the FACS & Imaging Core Facility for their help with microscopy; Ilian Atanassov and the Proteomics Core Facility for their help with proteomics data; Patrick Giavalisco and Yvonne Hinze from the Metabolomics Core Facility for their help with UPLC-MS; Jorge Boucas and the Bioinformatics Core Facility for their help with analysis of RNA-seq data. We thank Astrid Schauss and Beatrix Martini from the Imaging Core Facility at CECAD for their support in generating the electron microscopy data. We gratefully acknowledge Andrea Hartmann, Sandra Buschbaum, André Pahl, Ramona Jansen and the rest of Mouse Tissue Bank team for help with mouse dissections and Oliver Hendrich for organizational assistance. The research leading to these results has received funding from the European Research Council under the European Union’s Seventh Framework Programme (FP7/2007–2013)/ ERC grant agreement no. 268739 and the European Union’s Horizon 2020 research and innovation programme no. 741989. We gratefully acknowledge funding from the Max Planck Society.

## Author contributions

P.J. and L.P. conceptualized study, P.J., S.G., Y.X.L., T.L. and L.P. designed the experiments, P.J., Y.X.L., T.L., L.F.D., J.L., T.N., S.A., J.C.R., E.F. and J. F. conducted the experiments, P.J., Y.X.L., T.L. analyzed the data, J.P. analyzed the proteomics data, P.J., S.G. and L.P. wrote original manuscript, P.J., Y.X.L., S.G. and L.P. edited it, P.J., Y.X.L. and T.L. contributed equally.

