## Extended Data for "Profound geroprotection from brief rapamycin treatment in early adulthood by persistently increased intestinal autophagy"

**­Extended Data Figures and Tables**

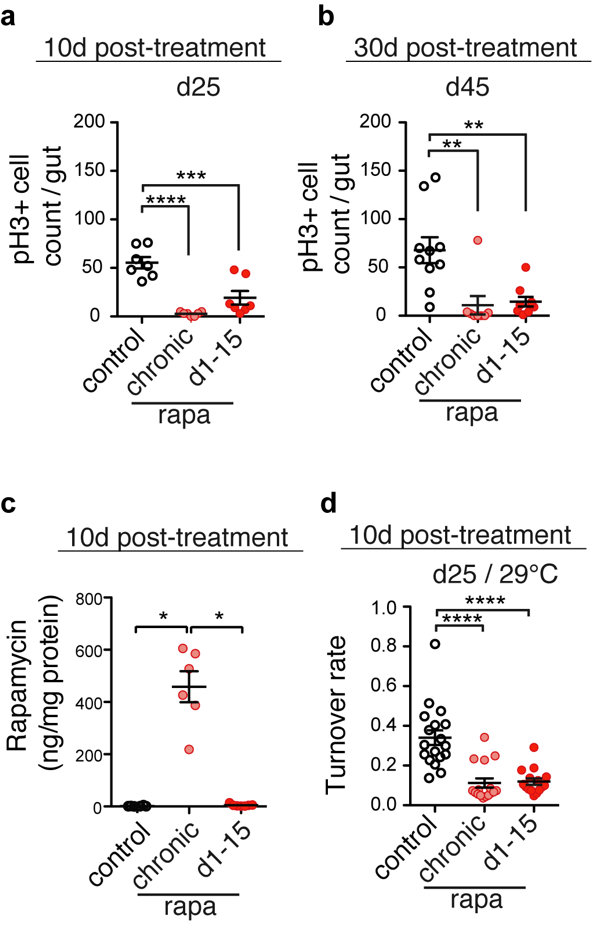

Extended Data Fig. 1. Brief rapamycin treatment inhibits ISC division and epithelial turnover rate to the late stages of life. a-b, Rapamycin treatment from day 1-15 reduced pH3+ cell number 10 (a) and 30 (b) days post-treatment to the same degree as rapamycin chronic exposure. c, Rapamycin concentration was increased in the guts of 25-day-old flies chronically treated with rapamycin but was absent from the intestine 10-days after rapamycin treatment was terminated. n=6-7. d, Turnover of intestinal epithelium was reduced 10 days after the treatment termination as assessed by esg^ts^ F/O system activation (at 29°C). Data are mean ± s.e.m. One-way ANOVA followed by Bonferroni’s post-test. **p < 0.01, ***p < 0.001, ****p < 0.0001.

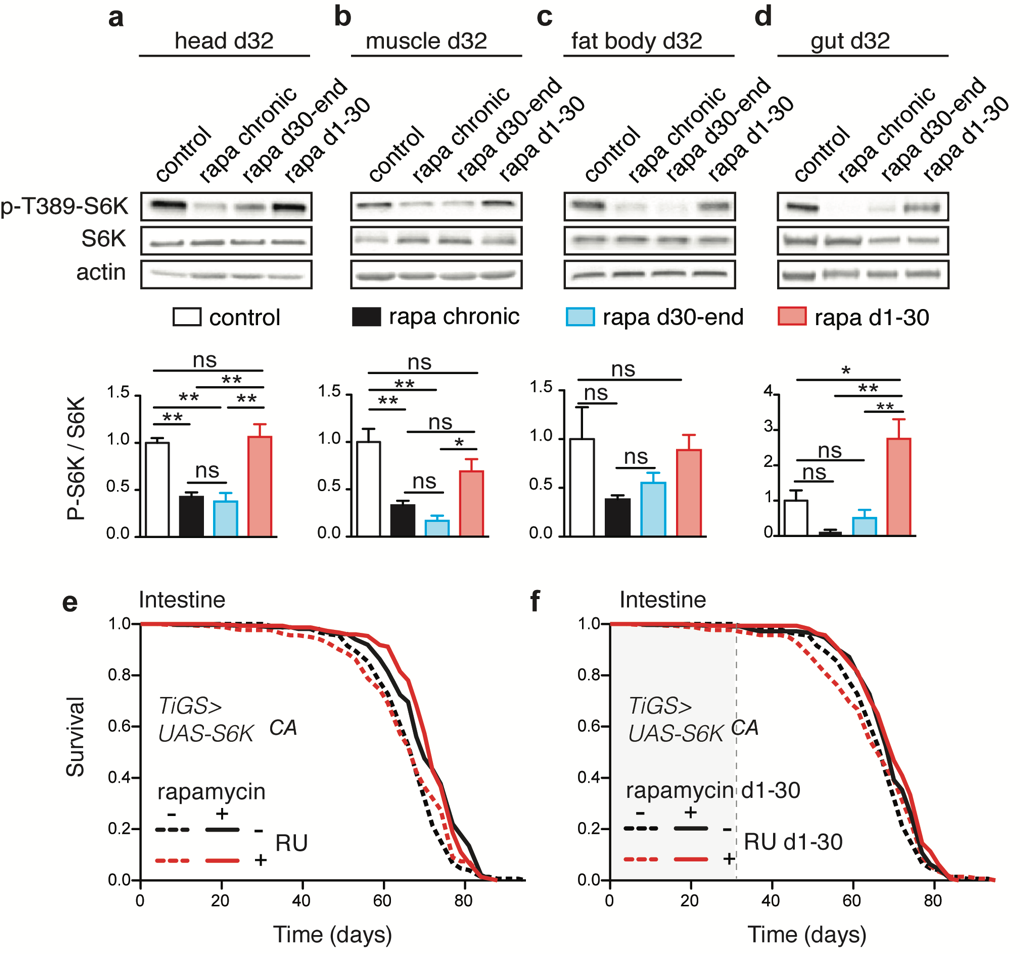

Extended Data Fig. 2. Effect of short-term rapamycin treatment on tissue-specific TORC1 activity and lifespan of flies over-expressing constitutively active S6K in the gut. a-d, Rapamycin treatment initiated on day 30 down-regulated TORC1 activity within 48h and terminating rapamycin treatment on day 30 de-repressed TORC1 activity back to the levels of control flies within 48h in heads, muscle, fat body and gut. Data are mean ± s.e.m. One-way ANOVA; Tukey’s multiple comparison test. N=3. *p < 0.05, **p < 0.01, ***p < 0.001. e-f, Chronic (p=9.95 x 10^-05^) and short-term (p=0.029) rapamycin treatment extended lifespan of control flies and of flies over-expressing constitutively active S6K in the gut (chronic: p=0.002, short-term treatment: p=0.02, see also Extended Data Table 4). N=200 flies per experimental group. Log-rank test and CPH analysis.

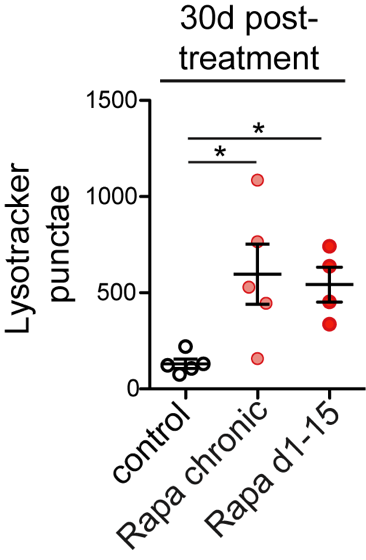

Extended Data Fig. 3. Effect of brief rapamycin treatment on the number of lysotracker+ punctae in the gut 30-days post-treatment. Rapamycin induced lysotracker-stained punctae in the gut of 45-days old flies, and these stayed induced 30-days post-treatment. Data are mean ± s.e.m. One-way ANOVA; Bonferroni’s multiple comparison test. *p < 0.05.

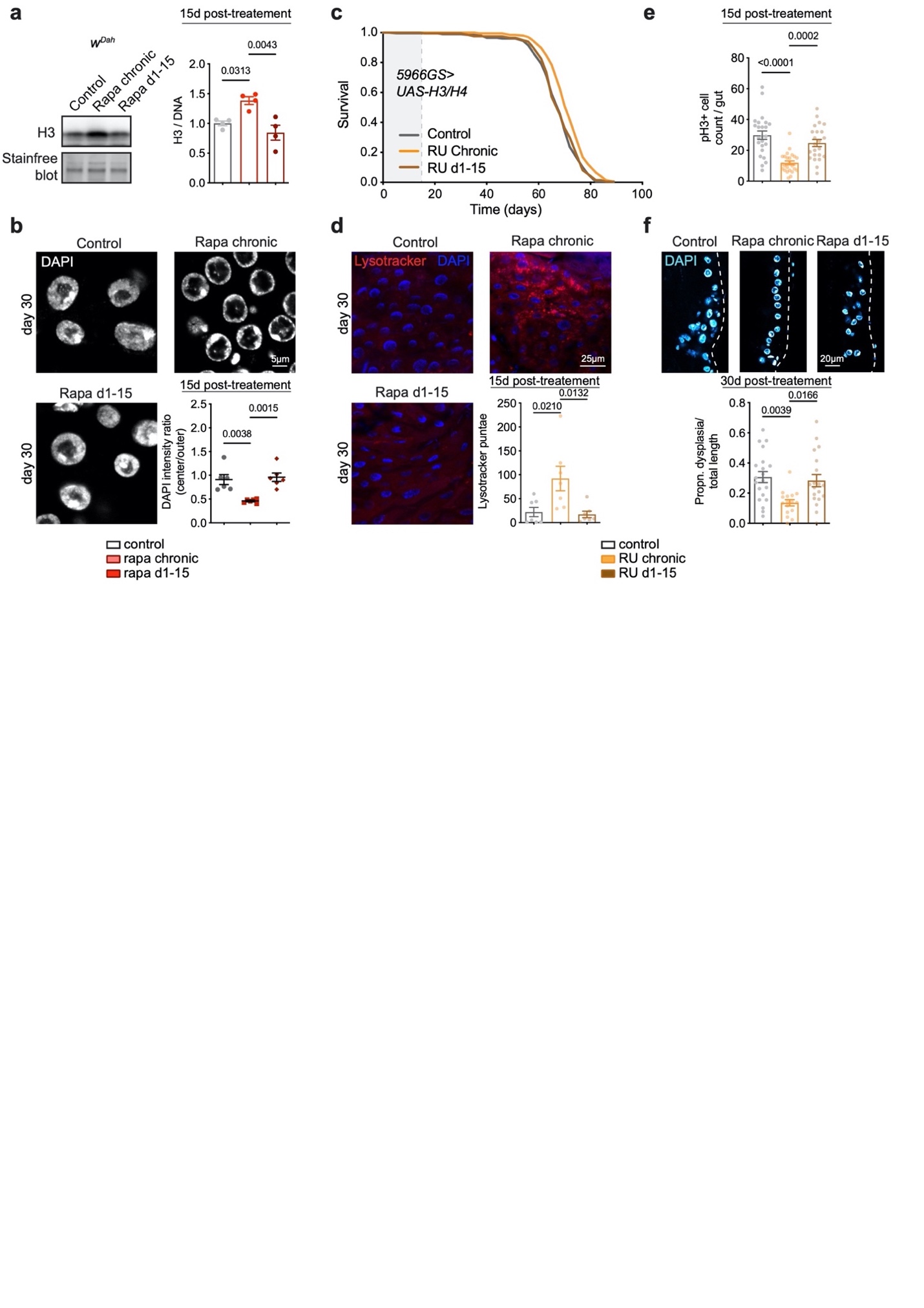

**Extended Data** **Fig. 4. Increased expression of histones by chronic rapamycin treatment is not responsible for the effects of short-term rapamycin treatment on lifespan and gut health.**

**a**, Immunoblot of histone H3 in the fly gut on day 30, 15-days post-rapamycin-treatment (n=4). **b**, The accumulation of chromatin at the nuclear envelope in ECs disappeared 15 days after d1-15 treatment was terminated (n=6 guts). **c**, Chronic but not d1-15 over-expression of histones H3/H4 specifically in enterocytes extended lifespan (chronic: p=7.39 x10^-06^; d1-15 treatment: p=0.60. n=160-200 flies per experimental group. Log-rank test. **d,** Number of punctae stained by LysoTracker in 30-day old flies overexpressing H3/H4 specifically in enterocytes chronically or in days 1-15 (n=7). **e,** The number of pH3+ cells in the gut 30 days after short-term rapamycin treatment was withdrawn (n=23-25). **f,** Intestinal dysplasia in gut R2 region of flies 30-days after short-term rapamycin treatment was terminated (n=16-20). Data are mean ± s.e.m. One-way ANOVA; Bonferroni’s multiple comparison test.

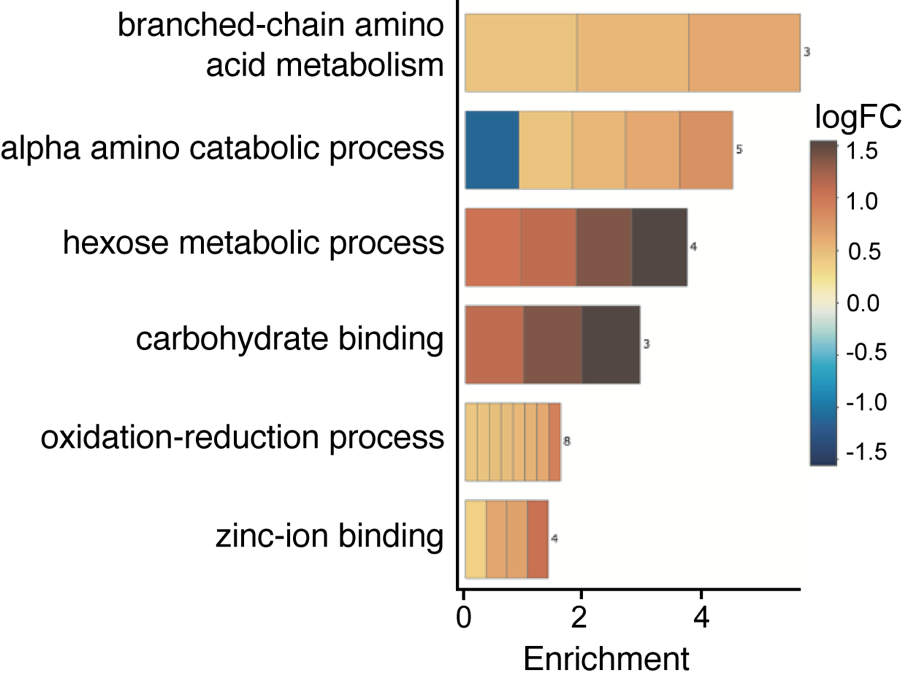

Extended Data Fig. 5. Rapamycin induces lasting changes in proteome of the gut. GO term analysis of proteins enriched by rapamycin treatment on day 25 that were also enriched 10 days after the drug was withdrawn.

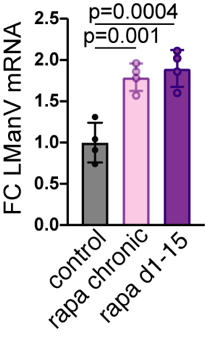

**Extended Data** **Fig. 6. Intestinal LManV mRNA levels.** Rapamycin treatment increased intestinal LManV mRNA, which stayed increased 10-days after the treatment was terminated. Data are mean ± s.e.m. One-way ANOVA; Dunnett’s multiple comparison test.

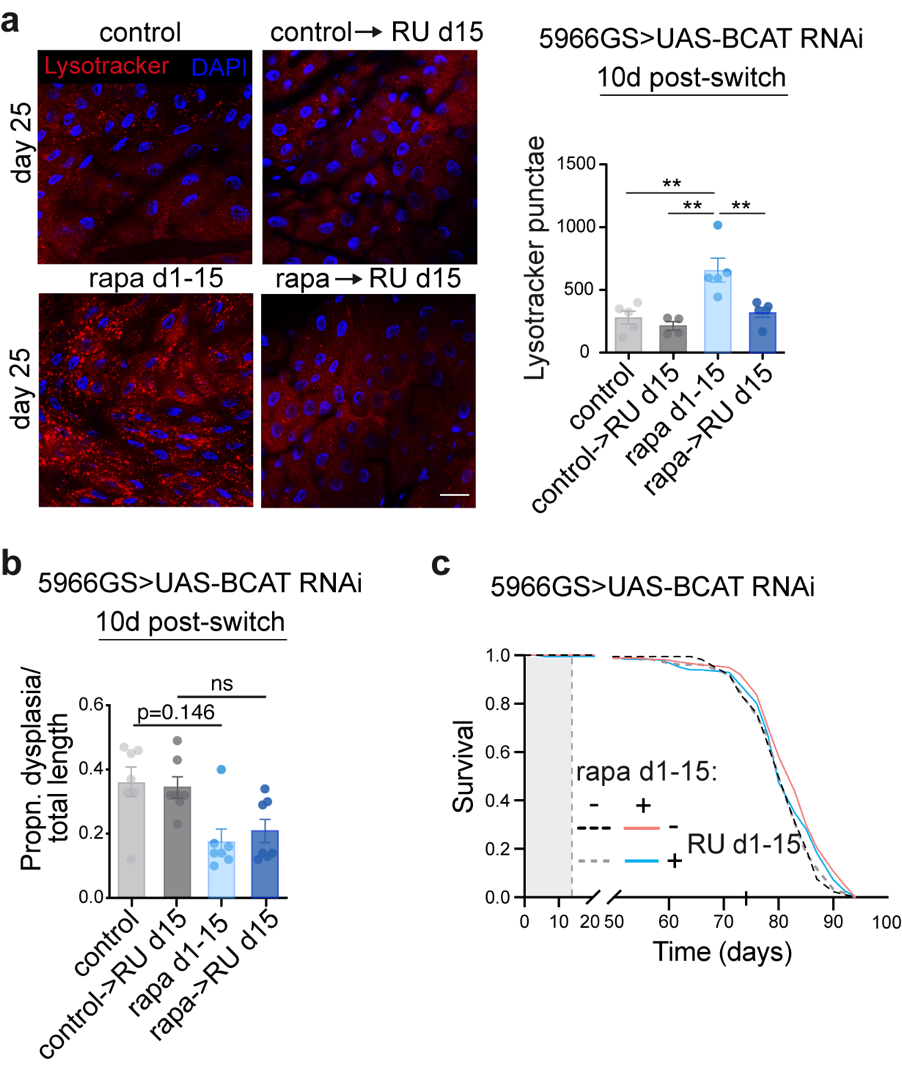

**Extended Data** **Fig.7. Persistent increase in BCAT mediates the ‘memory of autophagy’, and partially mediates gut pathology and lifespan induced by short-term rapamycin treatment.** **a-b**, Number of LysoTracker stained punctae in intestines (**a**) and gut pathology (**b**) of flies treated with rapamycin from day 1-15 followed by over-expression of RNAi against BCAT (n=5-7). Data are mean ± s.e.m. Two-way ANOVA; Bonferroni’s multiple comparison test. **p < 0.01. ***p < 0.001. **c**, Brief rapamycin (p=7.8 x 10^-4^) treatment extended lifespan of control flies, but not of flies expressing RNAi against BCAT in enterocytes (p=0.14, see also Extended Data Table 9). Log-rank test. n=180.

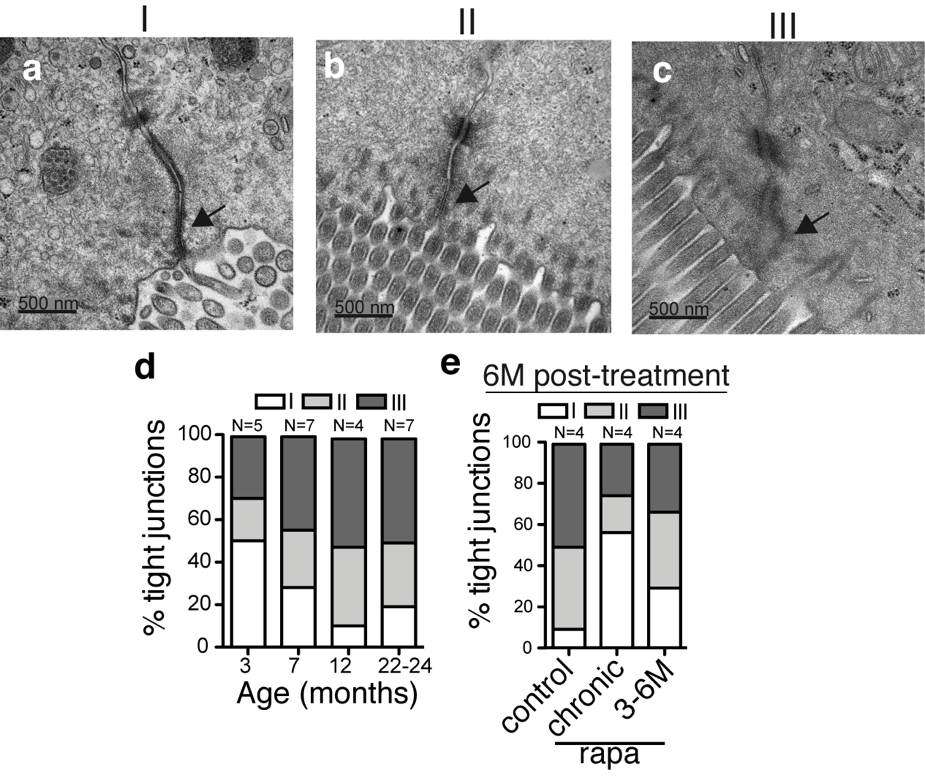

Extended Data Fig. 8. Effects of ageing and chronic and short-term rapamycin treatment on TJ. a-c, Tight junction pathology score: I - narrow and electron dense TJs (a); II - reduced electron density, but no dilations within TJs (b); III - low electron density and dilated TJs (c). d, Number of compromised TJs increased by 7 months of age compared to 3 months old mice (3M v. 7M: p<0.05; 3M v. 12M: p<0.0001; 3M v. 22-24M: p<0.0001; 7M v. 12M: ns; 12M v. 22-24M: ns). e, Rapamycin reduced the percentage of compromised TJs (class II and III) in mouse jejunum. Numbers of mice are indicated above each bar; at least 10 TJs per mouse were analyzed. One-way ANOVA followed by Bonferroni’s test performed using the number of intact TJs. One-way ANOVA followed by Bonferroni’s test. *p < 0.05, ***p < 0.001.

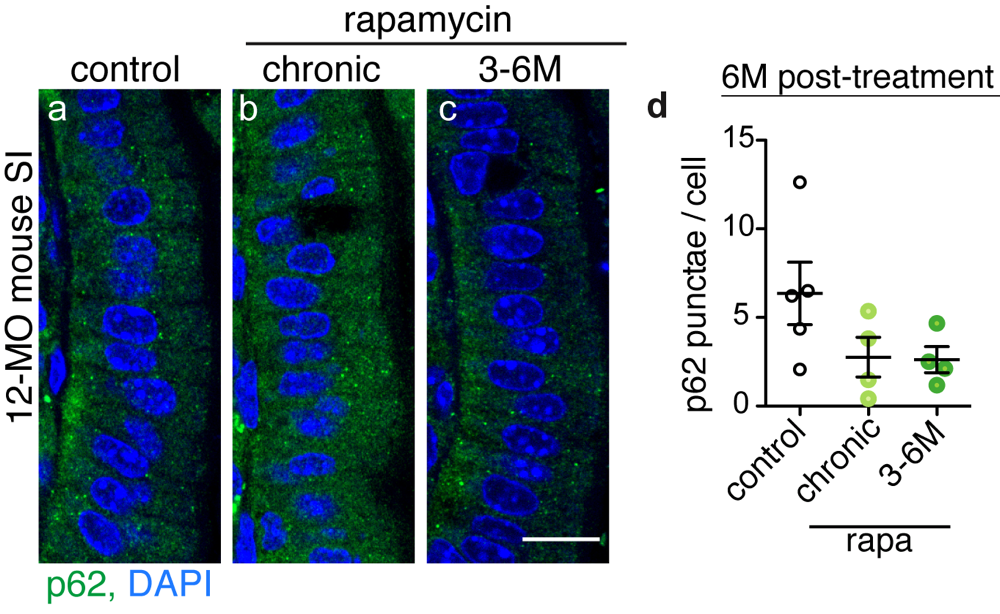

Extended Data Fig. 9. Rapamycin does not significantly affect the number of p62 punctae in the villi region of small intestine. n=4-5 mice, 50 or more cells in 5 different villi per mouse were analyzed. Data are mean ± s.e.m. One-way ANOVA, Bonferroni’s post-test. Scale is 15 μm.

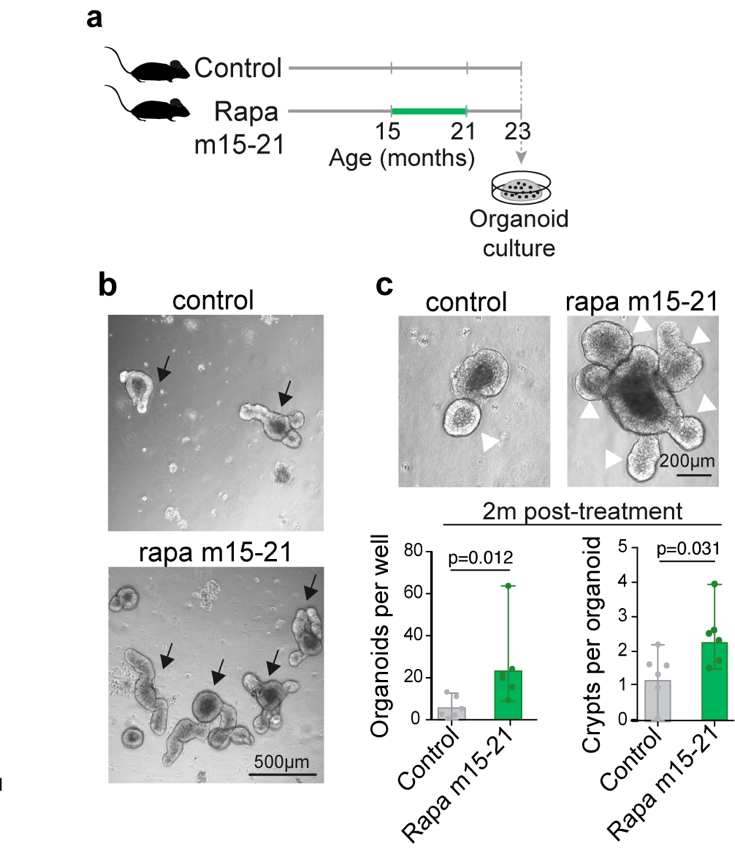

Extended Data Fig. 10. Brief rapamycin treatment improves regenerative capacity of the mammalian intestinal epithelium. a, Experimental design. b-c, Rapamycin increased organoid forming potential of intestinal crypts (arrows) and (j), formation of de novo crypts (white arrowheads), which remained increased 2 months post-treatment (n=6). Data are mean ± s.d. Two-sided student t-test.

Extended Data Table 1. Statistical analysis of survival data presented in Fig. 1a-d.

| n=400/treatment | | | |
| --- | --- | --- | --- |
| **Rapamycin day 45-end** – data before day 45 were censored | | | |
| **p-value (log rank)** |  |  |  |
|  | control | rapamycin | rapamycin 45-end |
| median (d) | 75.5 | 82 | 78 |
| mean (d) | 73.8 | 83.2 | 76.3 |
| control |  | 1.22E-28 | 0.00030 |
| rapamycin chronic |  |  | 3.7E-12 |
| **Rapamycin day 60-end** – data before day 60 were censored | | | |
| **p-value (log rank)** |  |  |  |
|  | control | rapamycin | rapamycin 60-end |
| median (d) | 75.5 | 84 | 75.5 |
| mean (d) | 76.9 | 84 | 77.7 |
| control |  | 1.09E-21 | 0.256315 |
| rapamycin chronic |  |  | 5.38E-16 |
| **Cox Proportional Hazard (CPH) analysis** **Parameter Estimates** \| Term \| \| Estimate \| \| SE \| \| p \| \|; \| rapamycin \| \| 0.13680 \| \| 0.025308 \| \| <0.0001 \| \|; d30 v d45 start \| timing \| \| 0.00355 \| \| 0.024808 \| \| 0.8861 \| \|; \| rapa*timing \| \| 0.00975 \| \| 0.024808 \| \| 0.6943 \| \|; \| rapamycin \| \| 0.07538 \| \| 0.027004 \| \| 0.0052 \| \|; d45 v d60 start \| timing \| \| 0.05899 \| \| 0.026677 \| \| 0.0267 \| \|; \| rapa*timing \| \| 0.08447 \| \| 0.026677 \| \| 0.0016 \| \| | | |  |

Extended Data Table 2. Statistical analysis of survival data presented in Fig. 1a and 1d.

| 30-day switch – data before day 30 were censored | | | | | |
| --- | --- | --- | --- | --- | --- |
| n=400/treatment |  |  |  |  |  |
| **p-value (log rank)** |  |  |  |  |  |
|  | control | rapamycin | rapamycin 30-end | rapamycin 1-30 |  |
| median (d) | 72.5 | 82 | 78 | 82 |  |
| mean (d) | 73.2 | 82.3 | 77.6 | 81.3 |  |
| control |  | 6.84E-27 | 2.13E-06 | 2.39E-13 |  |
| rapamycin chronic |  |  | 1.04E-13 | 0.087215 |  |
| rapamycin 30-end |  |  |  | 2.88E-06 |  |
| **Cox Proportional Hazard (CPH) analysis** | | |  |  |  |
| **Risk Ratios** |  |  |  |  |  |
| Level 1 | /Level 2 | Risk Ratio | p | Lower 95% | Upper 95% |
| rapamycin | control | 0.4309059 | <0.0001 | 0.3734110 | 0.4977745 |
| rapamycin 1-30 | rapamycin chronic | 1.1543417 | 0.1266 | 0.9601738 | 1.3877745 |
| rapamycin 30-end | rapamycin chronic | 1.4612031 | <0.0001 | 1.2173019 | 1.7539727 |
| rapamycin 1-30 | rapamycin 30-end | 0.6843676 | <0.0001 | 0.5701343 | 0.8214889 |

Extended Data Table 3. Statistical analysis of survival data presented in Fig. 1e and 2a.

| 15-day switch |  |  |  |  |  |
| --- | --- | --- | --- | --- | --- |
| n=400/treatment |  |  |  |  |  |
| **p-value (log rank)** |  |  |  |  |  |
|  | control | rapamycin | rapamycin 1-30 | rapamycin 15-30 | rapamycin 1-15 |
| median (d) | 72.5 | 77.5 | 80 | 77.5 | 77.5 |
| mean (d) | 72.1 | 77.7 | 77.5 | 76.0 | 77.5 |
| maximum (d) | 89 | 91.5 | 91.5 | 91.5 | 91.5 |
| control |  | 1.2944E-10 | 2.0562E-13 | 7.585E-07 | 1.2016E-11 |
| rapamycin chronic |  |  | 0.14736386 | 0.1908386 | 0.50923849 |
| rapamycin 1-30 |  |  |  | 0.0153711 | 0.51293879 |
| rapamycin 15-30 |  |  |  |  | 0.07282181 |
| **Cox Proportional Hazard (CPH) analysis** | | |  |  |  |
| **Risk Ratios** |  |  |  |  |  |
| Level 1 | /Level 2 | Risk Ratio | p | Lower 95% | Upper 95% |
| rapamycin | control | 0.6522501 | <0.0001 | 0.565943 | 0.751711 |
| rapamycin 1-15 | rapamycin chronic | 0.9632981 | 0.6074 | 0.834742 | 1.111281 |
| rapamycin 15-30 | rapamycin chronic | 1.0933126 | 0.2117 | 0.950452 | 1.257805 |
| rapamycin 1-15 | rapamycin 15-30 | 0.8809997 | 0.0792 | 0.764565 | 1.014886 |

Extended Data Table 4. Statistical analysis of survival data presented in Extended Data Fig. 2e-f.

| *TiGS/UAS-S6K const.act.* | |  |  |  |  |  |  |
| --- | --- | --- | --- | --- | --- | --- | --- |
| n=200/treatment | |  |  |  |  |  |  |
| **p-value (log rank)** | |  |  |  |  |  |  |
|  | EtOH | RU | Rapa | Rapa + RU | RU 1-30 | Rapa 1-30 | Rapa + RU 1-30 |
| median (d) | 67 | 67 | 69 | 71 | 65 | 69 | 69 |
| mean (d) | 66.5 | 65.8 | 70.4 | 71.4 | 63.8 | 67.3 | 69.6 |
| maximum (d) | 75.5 | 75.5 | 82.5 | 82.5 | 75.5 | 80 | 82.5 |
| EtOH |  | 0.37293 | 9.9E-05 | 9.27E-06 | 0.424 | 0.029619 | 0.001376 |
| RU |  |  | 0.00438 | 0.002346 | 0.8139 | 0.396167 | 0.071006 |
| Rapamycin |  |  |  | 0.863832 | 0.0017 | 0.016441 | 0.16678 |
| Rapamycin + RU |  |  |  |  | 0.0013 | 0.011603 | 0.229435 |
| RU 1-30 |  |  |  |  |  | 0.225444 | 0.020668 |
| Rapamycin 1-30 |  |  |  |  |  |  | 0.23175 |
| **Cox Proportional Hazard (CPH) analysis** | | | |  |  |  |  |
| **Risk Ratios** |  |  |  |  |  |  |  |
| Level 1 | /Level 2 | Risk Ratio | p | Lower 95% | Upper 95% |  |  |
| rapa chronic | control | 0.69202 | <0.0001 | 0.587187 | 0.8156 |  |  |
| induced chronic | non-induced chronic | 0.94363 | 0.4864 | 0.801264 | 1.1112 |  |  |
| rapa 1-30 | control | 0.0799 | 0.0064 | 0.680384 | 0.9388 |  |  |
| induced 1-30 | non-induced 1-30 | 0.90581 | 0.2265 | 0.771479 | 1.0633 |  |  |
| **Parameter Estimates** | |  |  |  |  |  |  |
|  | Term | Estimate | SE | p |  |  |  |
| chronic | rapamycin | 0.18407 | 0.0418 | <0.0001 |  |  |  |
|  | induction | -0.0290 | 0.0416 | 0.4864 |  |  |  |
|  | rapamycin*induction | -0.0271 | 0.0418 | 0.5164 |  |  |  |
| day 1-30 | rapamycin | 0.11219 | 0.0410 | 0.0064 |  |  |  |
|  | induction | -0.0494 | 0.0400 | 0.2265 |  |  |  |
|  | rapamycin*induction | -0.0045 | 0.0408 | 0.911 |  |  |  |

| **p-value (log rank)** | | | | |  | |  | |  | |  |  |  |
| --- | --- | --- | --- | --- | --- | --- | --- | --- | --- | --- | --- | --- | --- |
|  | | EtOH | | | RU | | Rapa | | Rapa + RU | | RU 1-15 | Rapa1-15 | Rapa+ RU 1-15 |
| median (d) | | 57 | | | 59 | | 61.5 | | 61.5 | | 61.5 | 64 | 61.5 |
| mean (d) | | 58.2 | | | 59.1 | | 62.7 | | 59.9 | | 59.1 | 63.7 | 60.9 |
| maximum (d) | | 68 | | | 70.5 | | 70.5 | | 68 | | 70.5 | 70.5 | 70.5 |
| EtOH | |  | | | 0.467679 | | 1.53E-05 | | 0.156857 | | 0.43617 | 8.8E-10 | 0.012342 |
| RU | |  | | |  | | 7.48E-05 | | 0.253446 | | 0.84502 | 2.4E-08 | 0.056321 |
| rapamycin | |  | | |  | |  | | 0.000263 | | 9.8E-05 | 0.03165 | 0.030291 |
| rapamycin + RU | |  | | |  | |  | |  | | 0.51559 | 5.4E-09 | 0.327552 |
| RU 1-15 | |  | | |  | |  | |  | |  | 7.4E-09 | 0.097373 |
| rapamycin 1-15 | |  | | |  | |  | |  | |  |  | 8.44E-05 |
| rapamycin + RU 1-15 | |  | | |  | |  | |  | |  |  |  |
| **Cox Proportional Hazard (CPH) analysis** | | | | | | | | |  | |  |  |  |
| **Risk Ratios** | | | | |  | |  | |  | |  |  |  |
| Level 1 | | /Level 2 | | | Risk Ratio | | p | | Lower 95% | | Upper 95% |  |  |
| rapa chronic | | control | | | 0.740356 | | 0.0006 | | 0.624593 | | 0.87815 |  |  |
| induced chronic | | non-induced chronic | | | 1.123287 | | 0.1774 | | 0.948662 | | 1.33023 |  |  |
| rapa 1-15 | | control | | | 0.553101 | | <0.0001 | | 0.461037 | | 0.66278 |  |  |
| induced 1-15 | | non-induced 1-15 | | | 0.960025 | | 0.6559 | | 0.802404 | | 1.149 |  |  |
| **Parameter Estimates** | | | | |  | |  | |  | |  |  |  |
|  | | Term | | | Estimate | | SE | | p | |  |  |  |
| chronic | | rapamycin | | | 0.150312 | | 0.043435 | | 0.0006 | |  |  |  |
|  |  | induction | | | 0.05813 | | 0.043093 | | 0.11774 | |  |  |  |
|  |  | rapamycin*induction | | | -0.083593 | | 0.043034 | | 0.0516 | |  |  |  |
| day 1-15 | | rapamycin | | | 0.203859 | | 0.044964 | | <0.0001 | |  |  |  |
|  |  | induction | | | 0.0543 | | 0.044685 | | 0.2246 | |  |  |  |
|  |  | rapamycin*induction | | | -0.074401 | | 0.044552 | | 0.0948 | |  |  |  |

**Extended Data** **Table 5**. Statistical analysis of survival data present­ed in Fig. 4a-b.

| *5966GS/UAS-Atg5 ^RNAi^* |
| --- |
| n=400/treatment |

Extended Data Table 6. Statistical analysis of survival data presented in Fig. 4c-d.

| *5961GS/UAS-Atg5 ^RNAi^* | | |  |  |  |  |  |
| --- | --- | --- | --- | --- | --- | --- | --- |
| n=200/treatment | |  |  |  |  |  |  |
| **p-value (log rank)** | |  |  |  |  |  |  |
|  | EtOH | RU | Rapa | Rapa + RU | RU 1-15 | Rapa 1-15 | Rapa + RU 1-15 |
| median (d) | 54.5 | 54.5 | 61.5 | 61.5 | 54.5 | 61.5 | 61.5 |
| mean (d) | 54.1 | 55.4 | 59.7 | 60.3 | 55.3 | 62.2 | 62.4 |
| maximum (d) | 68 | 70.5 | 69 | 68 | 61.5 | 70.5 | 70.5 |
| EtOH |  | 0.233667 | 3.4E-06 | 3.49E-06 | 0.5718 | 8.2E-13 | 1.93E-12 |
| RU |  |  | 0.001097 | 0.001491 | 0.4706 | 2.8E-09 | 1.1E-08 |
| rapamycin |  |  |  | 0.73857 | 1E-05 | 0.00529 | 0.011967 |
| rapamycin + RU |  |  |  |  | 2.4E-05 | 0.00053 | 0.001288 |
| RU 1-15 |  |  |  |  |  | 8.2E-12 | 5.4E-12 |
| rapamycin 1-15 |  |  |  |  |  |  | 0.386465 |
| rapamycin + RU 1-15 |  |  |  |  |  |  |  |
| **Cox Proportional Hazard (CPH) analysis** | | | |  |  |  |  |
| **Risk Ratios** | |  |  |  |  |  |  |
| Level 1 | /Level 2 | Risk Ratio | p | Lower 95% | Upper 95% |  |  |
| rapa chronic | control | 0.630867 | <0.0001 | 0.526928 | 0.75437 |  |  |
| induced chronic | non-induced chronic | 0.929143 | 0.4213 | 0.776204 | 1.11124 |  |  |
| rapa 1-15 | control | 0.553101 | <0.0001 | 0.461037 | 0.66278 |  |  |
| induced 1-15 | non-induced 1-15 | 0.960025 | 0.6559 | 0.802404 | 1.149 |  |  |
| **Parameter Estimates** | |  |  |  |  |  |  |
|  | Term | Estimate | SE | p |  |  |  |
| chronic | rapamycin | 0.23033 | 0.045737 | <0.0001 |  |  |  |
|  | induction | -0.036746 | 0.045736 | 0.4213 |  |  |  |
|  | rapamycin*induction | -0.022517 | 0.045743 | 0.6225 |  |  |  |
| day 1-15 | rapamycin | 0.138892 | 0.03734 | 0.0002 |  |  |  |
|  | induction | 0.15442 | 0.037407 | <0.0001 |  |  |  |
|  | rapamycin*induction | -0.024520 | 0.03722 | 0.5101 |  |  |  |

**Extended Data** **Table 7**. Statistical analysis of survival data presented in Fig. 5b-c.

| *5966GS/UAS-Atg1 ^OE^* | |  |  |  |  |
| --- | --- | --- | --- | --- | --- |
| n=200/treatment (Fig. 5b) and 160/treatment (Fig. 5c) | | | | | |
| **p-value (log rank)** | |  |  |  |  |
|  | EtOH | RU | Rapa | Rapa+ RU |  |
| median (d) | 60 | 67 | 64.5 | 64.5 |  |
| mean (d) | 59.2 | 66.4 | 64.4 | 63.5 |  |
| maximum (d) | 71.5 | 71.5 | 71.5 | 71.5 |  |
| EtOH |  | 3.6E-11 | 5.68E-06 | 8.49E-05 |  |
| RU |  |  | 0.046318 | 0.004648 |  |
| rapamycin |  |  |  | 0.506055 |  |
| rapamycin + RU |  |  |  |  |  |
|  | EtOH | RU 1-15 | Rapa 1-15 | Rapa + RU 1-15 | |
| median (d) | 60.5 | 67.5 | 67.5 | 67.5 |  |
| mean (d) | 60.3 | 66.8 | 67.3 | 67.8 |  |
| maximum (d) | 74.5 | 74.5 | 74.5 | 74.5 |  |
| EtOH |  | 6.72E-5 | 1.00E-06 | 1.84E-07 |  |
| RU 1-15 |  |  | 0.371120 | 0.212741 |  |
| rapamycin 1-15 |  |  |  | 0.697781 |  |
| rapamycin + RU 1-15 |  |  |  |  |  |
| **Cox Proportional Hazard (CPH) analysis** | | | |  |  |
| **Risk Ratios** | |  |  |  |  |
| Level 1 | /Level 2 | Risk Ratio | p | Lower 95% | Upper 95% |
| rapa chronic | control | 0.567652 | <0.0001 | 0.461557 | 0.70218 |
| induced chronic | non-induced chronic | 0.380319 | <0.0001 | 0.307387 | 0.47305 |
| rapa 1-15 | control | 0.725947 | 0.0002 | 0.613941 | 0.85838 |
| induced 1-15 | non-induced 1-15 | 0.790212 | 0.0059 | 0.668278 | 0.93439 |
| **Parameter Estimates** | |  |  |  |  |
|  | Term | Estimate | SE | p |  |
| chronic | rapamycin | -0.283123 | 0.053447 | <0.0001 |  |
|  | induction | -0.483371 | 0.054918 | <0.0001 |  |
|  | rapamycin*induction | 0.156973 | 0.052663 | 0.0036 |  |
| day 1-15 | rapamycin | 0.160138 | 0.042750 | 0.0002 |  |
|  | induction | -0.117726 | 0.042755 | 0.0059 |  |
|  | rapamycin*induction | -0.107443 | 0.042741 | 0.0119 |  |

Extended Data Table 8. Statistical analysis of survival data presented in Extended Data Fig. 4c.

| *5966GS/UAS-H3/H4 ^OE^*  n=200/treatment | | | |
| --- | --- | --- | --- |
| **p-value (log rank)** |  |  |  |
|  | EtOH | RU | RU 1-15 |
| median (d) | 66.6 | 71.0 | 66.6 |
| mean (d) | 66.9 | 71.0 | 67.5 |
| maximum (d) | 82.0 | 89.0 | 86.0 |
| control |  | 7.38E-06 | 0.59709 |
| RU |  |  | 0.00011 |

**Extended Data** **Table 9**. Statistical analysis of survival data presented in Extended Data Fig. 7c.

| *5966GS/UAS-BCAT ^RNAi^* | |  |  |  |
| --- | --- | --- | --- | --- |
| n=180/treatment | | | | |
| **p-value (log rank)** | |  |  |  |
|  | EtOH | RU 1-15 | Rapa 1-15 | Rapa + RU 1-15 |
| median (d) | 79.0 | 79.0 | 81.5 | 79.0 |
| mean (d) | 80.6 | 80.5 | 82.2 | 80.9 |
| maximum (d) | 86.0 | 88.5 | 91.0 | 88.5 |
| EtOH |  | 0.699135 | 7.83E-04 | 0.050575 |
| RU 1-15 |  |  | 0.002971 | 0.138767 |
| rapamycin 1-15 |  |  |  | 0.150979 |
| rapamycin + RU 1-15 |  |  |  |  |
